# Hiplot: a comprehensive and easy-to-use web service boosting publication-ready biomedical data visualization

**DOI:** 10.1101/2022.03.16.484681

**Authors:** Jianfeng Li, Benben Miao, Shixiang Wang, Wei Dong, Houshi Xu, Chenchen Si, Wei Wang, Songqi Duan, Jiacheng Lou, Zhiwei Bao, Hailuan Zeng, Zengzeng Yang, Wenyan Cheng, Fei Zhao, Jianming Zeng, Xue-Song Liu, Renxie Wu, Yang Shen, Zhu Chen, Saijuan Chen, Mingjie Wang, Hiplot Consortium

## Abstract

Modern web techniques provide an unprecedented opportunity for leveraging complex biomedical data generating in clinical, omics, and mechanism experiments. Currently, the functions for carrying out publication-ready biomedical data visualization represent primary technical hurdles in the state-of-art omics-based web services, whereas the demand for visualization-based interactive data mining is ever-growing. Here, we propose an easy-to-use web service, Hiplot (https://hiplot.com.cn), equipping with comprehensive and interactive biomedical data visualization functions (230+) including basic statistics, multi-omics, regression, clustering, dimensional reduction, meta-analysis, survival analysis, risk modeling, etc. We used the demo and real datasets to demonstrate the usage workflow and the core functions of Hiplot. It permits users to conveniently and interactively complete a few specialized visualization tasks that previously could only be done by senior bioinformatics or biostatistics researchers. A modern web client with efficient user interfaces and interaction methods has been implemented based on the custom components library and the extensible plugin system. The versatile output can also be produced in different environments via using the cross-platform portable command-line interface (CLI) program, Hctl. A switchable view between the editable data table and the file uploader/path selection could facilitate data importing, previewing, and exporting, while the plumber-based response strategy significantly reduced the time costs for generating basic scientific graphics. Diversified layouts, themes/styles, and color palettes in this website allow users to create high-quality and publication-ready graphics. Researchers devoted to both life and data science may benefit from the emerging web service.

## Introduction

Exploration and mining of multidimensional biomedical data originating from experimental assays, omics studies, and clinical observations largely rely on modern graphics and statistics [1], such as statistical description/inference and disease diagnosis [2]. In addition, visualization-based data mining techniques play an increasingly critical role in further enhancing interpretability, reproducibility, and effectiveness of both hypothesis- and data-driven scientific research [3-6]. Over a decade ago, users could only use desktop applications with limited functions and scalability to perform daily visualization analysis of scientific data. Recently, web-based cloud applications with better scalability have therefore become one of the ideal options leveraging complex biomedical data for biologists and clinicians who lack programming skills [7-13]. Since the establishment of well-known bioinformatics cloud services, such as Galaxy and DNAnexus [14], a number of upstream data analysis tasks have been moderately simplified, such as sequence alignments, variant calling, epigenetic profiling, and other workflow-based pipelines.

However, the common downstream functions including publication-ready scientific graphics and interactive data mining, visualization analysis in particular, based on the tabular data are still quite lacking in these websites [11, 15, 16]. The well-known bioinformatics cloud platform, Galaxy, only provides limited dozens of visualization-based plugins and still lacks adequate optimization for those lightweight biomedical visualization tasks. The visualization module of St. Jude Children’s Research Hospital cloud portal offers 20 JavaScript-based plugins for the interactive cancer genomics visualizations, while the basic scientific graphics is blank [12, 17]. The imageGP merely developed 16 subfunctions for scientific graphics and analysis since 2017 [18]. A huge amount of work still needs to be done for diverse visualization demands, which requires joint efforts from the entire scientific community. In addition, complicated user interfaces and inefficient interactions have become the major negative factors for users skipping web-based tools with functions of biomedical data visualization. For example, it has been rarely supported in the existing web-based bioinformatics tools that could preview and edit the data in the online spreadsheet editor, like classic desktop commercial graphics software. The automatic arrangements of multiple graphics in publication layout, e.g., 4, 6, 9 items per page, also often were overlooked. Other explicit issues, such as untimely tasks output, inconvenient reproduction of parameters/results, and lacking the cross-platform and easy-to-use command-line program, may further prevent the web-based tools from being more widely used for conducting biomedical data visualization tasks.

To meet these challenges, we propose an emerging easy-to-use and scalable web service, Hiplot (https://hiplot.com.cn), and an interdisciplinary community focusing on creating interactive applications related to biomedical data visualization. Since October 2019, hundreds of interactive web plugins related to visualization-based data mining have been developed by the Hiplot collaborative group. The core subfunctions of Hiplot have been built based on the open-source and published methods, e.g. R base graphics, ggplot2, pheatmap, ComplexHeatmap [2], circlize [19], ggstatsplot, cola [20], Broad Gene Set Enrichment Analysis (GSEA) [21], clusterProfiler [22], DISCOVER [23], etc. It has covered the most common demands of biologists and clinicians for daily biomedical data visualization using concise user interfaces and efficient interaction methods, and tens of thousands of researchers have already been using our web services. We expect this toolkit could become a useful infrastructure of data visualization for a broad of researchers in biomedicine, life sciences, and data science.

## Implementation

### Concise and easy-to-use web interfaces and interactions

The modern UI development framework Vuetify.js (v2) using Vue.js (v2) syntax and in-house components library were integrated to construct the website. One of the major advantages of the Hiplot web service is its user-friendly and efficient web interfaces and interaction methods (**Fig. S1, S2, S3**). Based on the web development framework of Hiplot, we provide a comparatively uniform user experience in high-frequency modern scientific graphics. The plugins of this website share a similar web layout following two columns design (**Fig. S3**). The left column shows the thumbnail, data, parameters, task list, and output of plugins, while the right column contains all static content including the description and the documentation of plugins. Thus, the vast majority of user interactions can be done on a single working page.

On the other hand, the online editable spreadsheet in data importing steps has been first introduced into this large-scale biomedical visualization cloud service with hundreds of plugins, which has better readability and editability to process small chunks of tabular data (**Fig. S4**). Meanwhile, the combination strategy of spreadsheet tabular input and switchable file uploader could improve the importing efficiency for different sizes of data files (**Fig. S4**). We implemented all operations of the core functional buttons and the switchable file uploader mode in the “hiplot-table-editor” web component. It supports preview, import, and edit the data in the spreadsheet view, which was developed based on the x-spreadsheet project [24] and Vuetify.js framework. Besides, it has been wrapped as a Vue component for more easily integrating into the web plugins of Hiplot. Through binding the options using table headers, the unique values of rows, and columns, the interactivity of visualization plugins has been further enhanced.

The pdf-collage plugin allows users to freely combine multiple visualization graphics of Hiplot in publication-ready layouts in batches. Previously, users needed to complete the routine task in the Adobe illustrator program with massive mouse clicks and drag-and-drop (**Fig. S5**). A pool of themes and hundreds of color palettes provided in the web plugins permit users to get high-quality and publication-ready outputs. The object of R data (.Rdata) can be used to reproduce or change the output style of graphics in the local R programming environment.

In terms of reproducibility of data, parameters, and issues, users can conveniently reproduce the history data/parameters and result in the plugins of Hiplot based on the standard JSON data objects from the local file or the remote file manager (**Fig. S6, S7**). To view the logging and input/output for debugging the possible errors via a confidential task index, Hiplot would generate a random temporary code in the browser cache for the newly submitted task, which has better task anonymity and security compared with the permanent cloud-based storage. Several interactive web applications based on the R Shiny and Python Streamlite development framework also have been developed for resolving different user demands (**Fig. S8**) [13]. The reproduction of errors in the Shiny- and Streamlite-based plugins are still challenging. Through the feedback system built in the Hiplot website, all submitted issues would be automatically synchronized to GitHub for unified management of issues.

### Cross-platform command-line interface program

The cross-platform command-line interface program of Hiplot (Hctl) was developed based on the Golang programming language, which conferred great advantages in helping users to quickly visualize multiple datasets in different environments at the same time. We adopted the minimal design principles, providing two core subcommands *config* and *plot* in the Hctl. Users could use the subcommand *config* of Hctl to query the demo data and parameters stored in a single JSON file. To submit a demo task of the heatmap plugin, users just need to execute the command “hctl plot -p <user_custom_input_path>/params.json -t heatmap -o <user_custom_output_path>”. The result files will be automatically retrieved when the task is finished or be manually requested via a random task key.

### JSON-based plugin system for visualization tasks

To increase the scalability of the website, we developed a JavaScript Object Notation (JSON)-based plugin system dynamically deploying new web plugins and simultaneously supporting the functionality of the web client and command-line program. All standard native plugins of Hiplot comprise multiple components including documentation, JSON files, and one or more core scripts. The documentation shown on the web page is directly parsed from individual multilingual Markdown files, which also could be reused in the VuePress documentation system. The JSON files rendering the plugins of Hiplot include *Meta JSON, UI JSON*, and *Data JSON* (**Fig. S9A**). The application name, thumbnail, entry, version, short description, maintainer, contact information, citation, release date, updated date, and quality score were stored in the *Meta JSON*. In contrast, *UI JSON* was designed to organize available web components of main data tables and parameters using the standard layout of plugins. *Data JSON* contains the default parameters and demo data that are indispensable for the web client, command-line program, and backend service.

Apart from the custom “hiplot-table-editor” and “cloud-file” web components, other seven core components include combobox, autocomplete, text-field, slider, range-slider, switch, and color-picker components of Vuetify.js framework have been supported in the *UI JSON*. The structured description of web interfaces has been significantly simplified the development of web clients for most data visualization analysis tasks. For example, the AGFusion program for visualizing chemical fusion genes has been wrapped in the AGFusionWeb project requiring massive development work in the front-end building and backend service [25], while we only need to use less than 200 lines of codes for realizing the web interface and backend function if as the plugin of the Hiplot website.

To reduce redundant parameters setting and programming handling, task parameters of *Data JSON* were divided into two parts. The general parameters were used in the web components and backend process functions that are shared in multiple plugins (**Fig. S9B**). The extra parameters only were used to control the specific task steps of plugins. Besides, we have been developed a standalone program HiSub [26] to render the JSON-based web plugin of Hiplot from a structured R script storing meta description, front-end fields, and backend utility (**Fig. S9C)**.

### Backend service and hardware layer improve tasks efficiency and reproducibility

It is noted that we used personalized task response methods for different visualization tasks with varied time costs, one of them is suitable for time-sensitive tasks. For example, under most circumstances, the users would like to obtain results of most basic visualization tasks in one to a few seconds. To reduce the time costs of R-based web plugins, we adopted a plumber-based multi-users R task response strategy in this work, allowing basic statistics graphics with multiple time-intensive dependencies to be completed in seconds. After starting a plumber session, the dependencies packages and environments will be pre-loaded as the resource representational state transfer (RESTful) application programming interfaces (APIs), which can efficiently process the new user task request and execute the backend R functions of task plugins. In contrast, the computation of gene enrichment analysis based on the clusterProfiler R program [22], and other command-line programs usually takes a few minutes. Because the loading of dependencies only occupies a small portion of the program’s runtime, there is no significant change in the time cost.

The control of the runtime environment of Hiplot is provided by Conda, Singularity image, and renv. The virtualization based on the Singularity image was used to install the dependence software with complex requirements. The renv-based runtime allows the Hiplot cloud service to be independent of the system R package environment and provides a stable and consistent package runtime environment for analysis reproducibility. The website is daily rolling updates in the development cycle. When the stable version is released, a snapshot of front-end *UI JSON*, background scripts, and third-party dependencies would be created. So, users can switch the version of Hiplot by using the selection button at the left top of the navigation.

In the hardware layer, two high-performance computing nodes with a high-speed internal network were included: 10 core CPUs with 40 threads, 90 TB storage, 128 GB memory; 20 core CPUs with 80 threads, 512 GB memory. Two computational nodes have been introduced for balancing task load and reducing network blockage. It is adequate to handle most lightweight tasks of biomedical data visualization based on tabular data.

## Results

### Overview of the comprehensive functions in the Hiplot

Since 2019, massive interactive web-based visualization applications (230+) have been developed by the Hiplot Consortium for different biomedical data mining tasks (**Fig. 1A, Table S1**). To our knowledge, this is one of the largest community-driven efforts to establish a free web service for interactively and comprehensively conducting publication-ready biomedical data visualization. In fact, the most of known modern statistics graphics have been implemented in this web service, which is comparable to the GraphPad, a well-known commercial desktop-based software for scientific data visualization (**Fig. 1B, C**). Hence, users can use these open accessed visualization tools of Hiplot to handle daily data analysis without limitations of the operating system or software environment for displaying and/or inferencing the data correlation, distribution, percentage, evolution, flow, ranking, and spatial features.

**Fig. 1.**
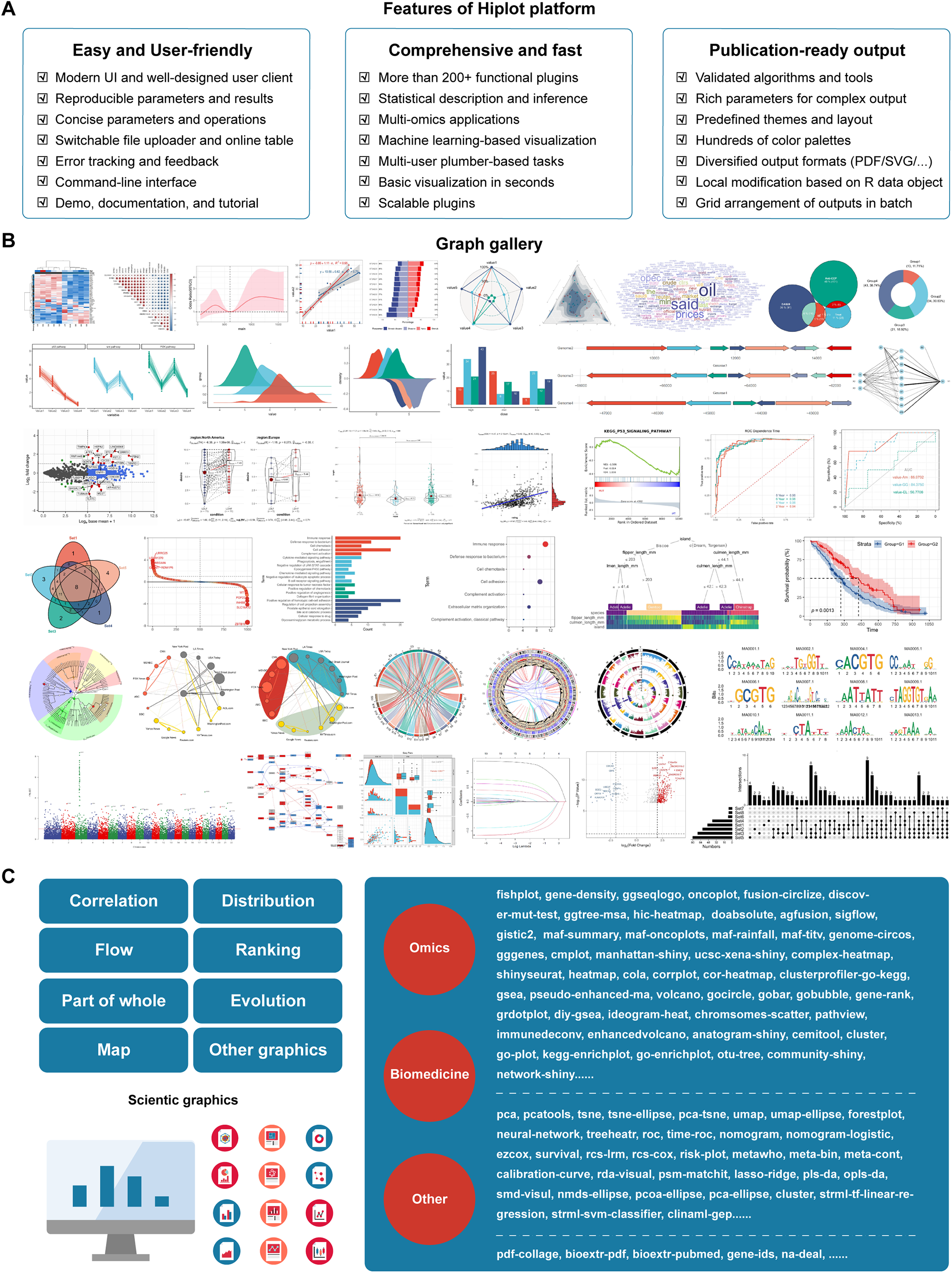
Overview of core features and functions in the Hiplot web service. (A) Three columns indicate the key features and advantages of the Hiplot cloud service. Comprehensive biomedical visualization functions related to modern statistical graphics, omics, and clinical data visualization have been established in the Hiplot website with a user-friendly web and command-line user interface. The spreadsheet and switchable file uploader simplified data importing for web-based lightweight visualization tasks. Basic graphics are completed in seconds based on the plumber workers. Users can directly use the Hiplot to generate publication-ready visualization graphics. (B) Graph gally shows selected demo outputs of partial plugins of Hiplot related to basic graphics, omics, and clinical data visualization. More demo output can be directly viewed in the cards list of applications on the website. (C) Classification of plugins and the four functions classes including basic graphics, omics, clinical, and other plugins. The left shows the core classes of basic graphics, and the right list the entry name of web plugins that were contributed by the Hiplot Consortium.

Excepting basic scientific graphics, users can freely and interactively explore cancer multi-omics datasets and comprehensively conduct multi-omics data visualizations, such as genome structure, chromosome distribution, genetic variations, population genetics, gene expression profiles, gene pathways enrichment, and tumor microenvironment (TME) (**Fig. 1B, C**). Besides, several machine learning-based visualization methods, such as unsupervised clustering, dimensional reduction algorithm (DRA), linear/non-linear regression, meta-analysis, survival analysis, and risk models, permit users to correlate multidimensional clinical features and conduct translational research (**Fig. 1B, C**).

### Comparison between Hiplot and similar visualization services

To demonstrate the advantages of Hiplot, we compared the Hiplot with representative websites (**Table 1**). The number and diversity of interactive visualization applications of Hiplot (native Hiplot, R Shiny, and Python Stremlit) are superior to other similar websites, the basic scientific graphics in particular. The implementation of many basic visualizations in Hiplot has attracted users from the whole scientific community, while the omics and clinical data visualizations may further assist the researchers of biomedicine and biology.

**Table 1.**
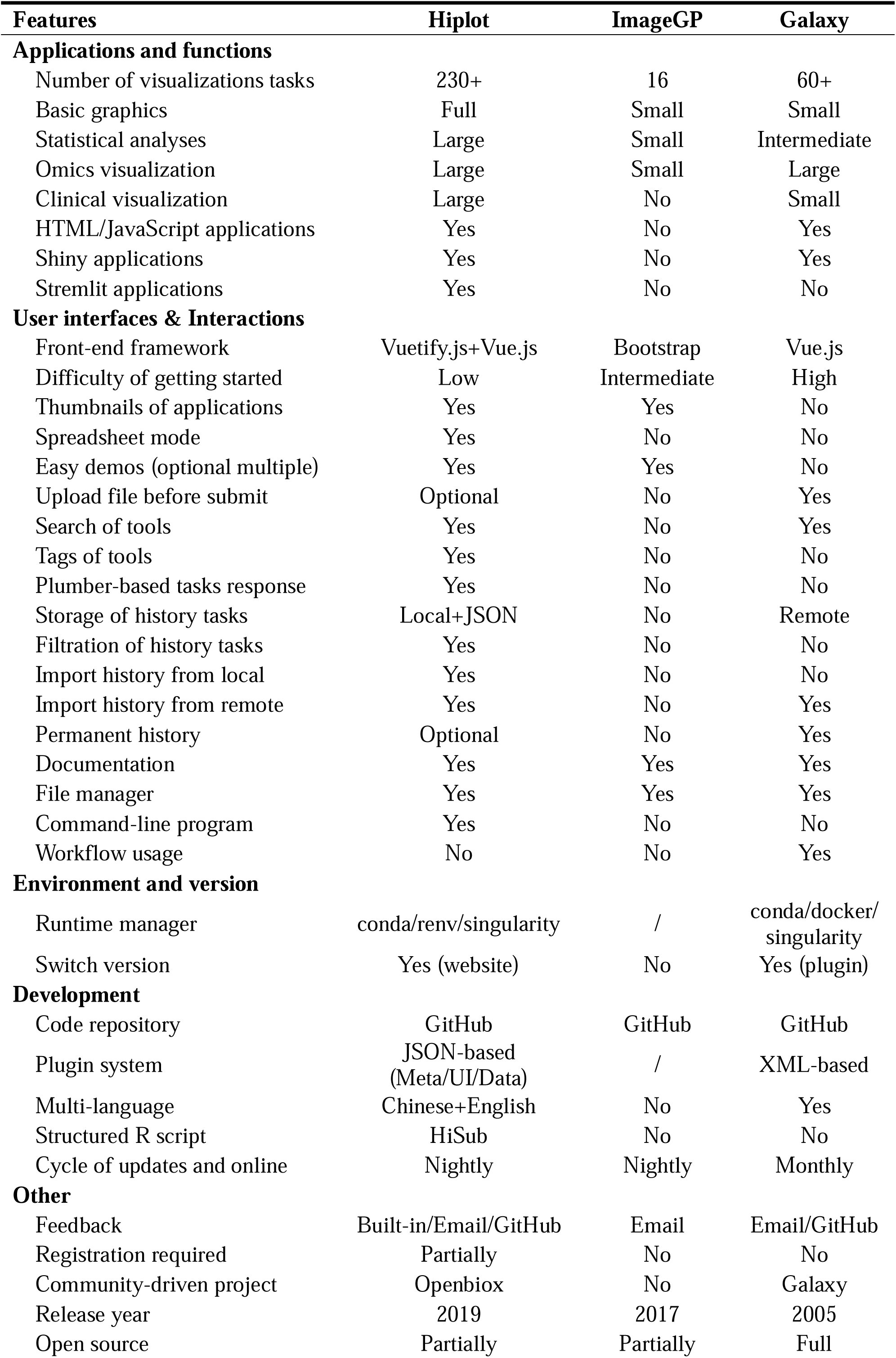
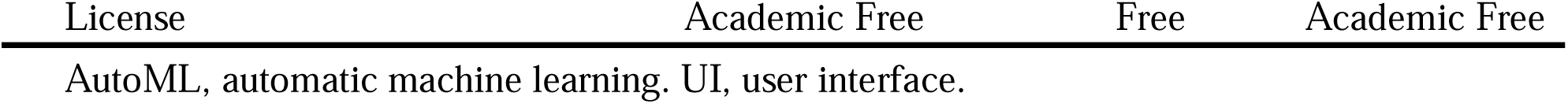
Comparison between Hiplot and similar web services.

In addition to the versatility of the task plugins, the efficient user interfaces and interaction methods in Hiplot, significantly save the learning cost of users. Users can invoke the plentiful functions of Hiplot via using the main web client or the command-line program (**Fig. 2A**). The web plugins in different modules, such as basic, advanced, clinical-tools, and mini-tools, can be quickly located via fuzzy matching of keyword and tag searching or switch and jump plugins via the top path. Thumbnails on plugins of cards allow users to find which web tools are needed. It is noted that in the traditional bioinformatics website, users only be allowed to upload the data using the file path selection or text area. The spreadsheet data editor with better readability and editability in the plugins of Hiplot may become an optional replacement method for other similar websites. Standard JSON data objects from local or remote storage can be used to reproduce the input and output of plugins in seconds. The multi-user plumber-based response method further improved the task efficiency in the lightweight visualization tasks (**Fig. 2B**) compared with the workflow-based response method requiring *de novo* loading dependencies. Though the setting up tasks using workflow editor has not been supported, it is feasible to integrate the functions of Hiplot into the existing data analysis pipeline via the command-line program. In the future, if there is a strong users demand, we can build a Galaxy-like interface using the existing plugins of Hiplot for better workflow-based integration.

**Fig. 2.**
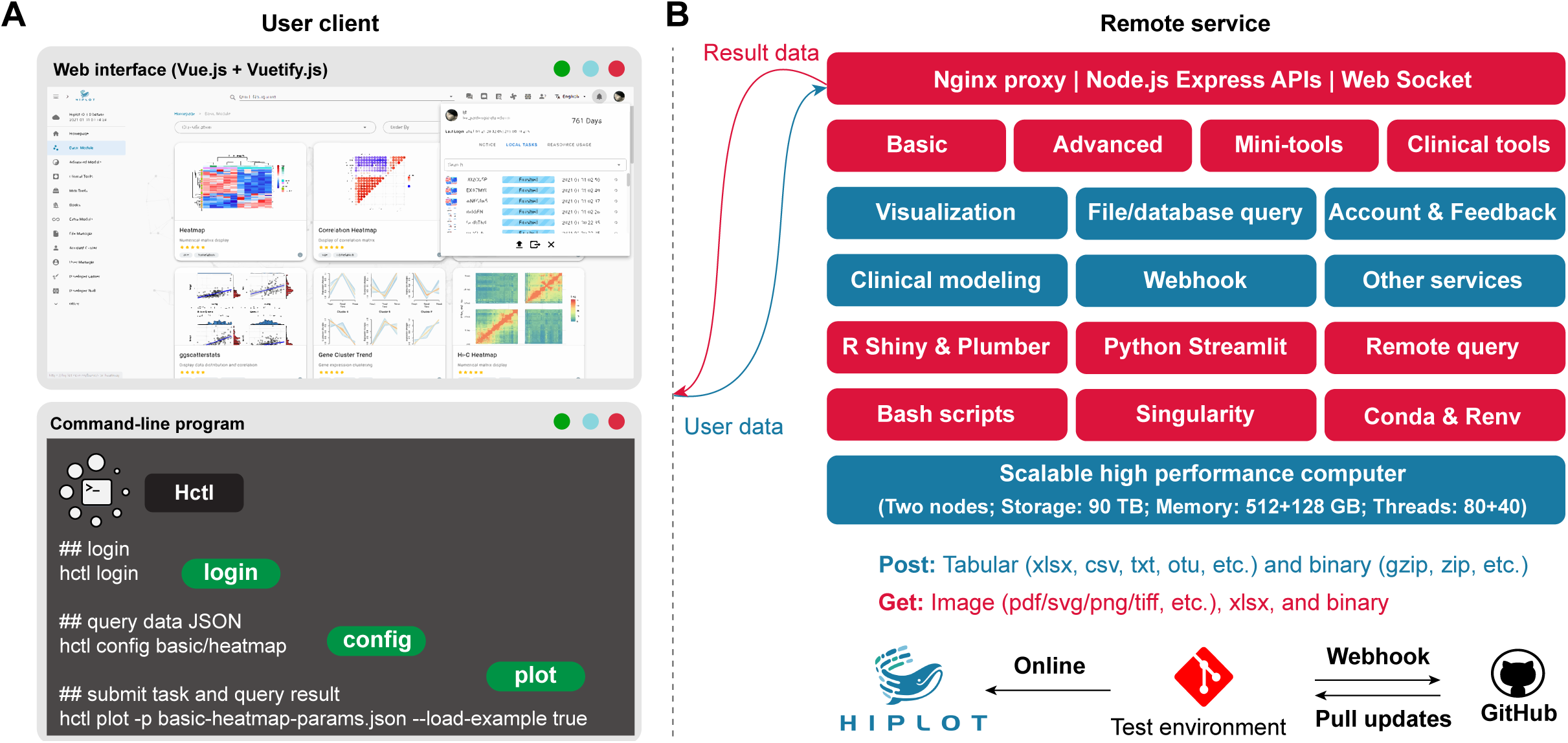
Website infrastructure and components from client to backend services. (A) Two types of user interfaces are provided including the web client and command-line program, Hctl. The top window shows the screenshot of the Hiplot web client on the basic module page and several cards of partial basic applications are listed. On the left of the page is the main menu to navigate the website in different modules. The notification window is also shown with the history task records. The bottom window shows the subcommands of Hctl including login, config, and submit. Login is required to use the Hctl program. The *config* and *submit* subcommands respectively to query the demo data/parameters and to submit tasks. (B) The infrastructure diagram illustrates the core backend services and hardware resources of the Hiplot web service. The web and command-line clients of Hiplot are communicated with the Nginx proxy/Node.js Express APIs/Web Socket services. The task plugins of Hiplot are distributed in four core modules including basic, advanced, mini-tools, and clinical tools. Apart from the JSON-based Vue.js plugin, the R Shiny and Python Streamlit frameworks are also introduced for the construction of interactive applications. The runtime environment of Hiplot was controlled by the renv, conda, and singularity. The plugins that are deployed in the Github of Hiplot or Openbiox organization will be automatically synced to the development environment or production environment if they pushed a new commit.

### Usage statistics represent popularity and potential impact

The project of Hiplot was launched in 2019, and the first version was released in March 2021. Here, we summarized the visits statistics of the Hiplot website from 09 July 2020 to 31 December 2021. It is encouraging that the Hiplot website has been visited more than 2,500,000 times from 100 countries worldwide in the web browser. Meanwhile, the website has reached averages of over 5,000 visits and 3,000 task submissions per day **(Fig. 3A**). More than 22,000 user accounts have been registered (**Fig. 3B**) and more than 160 plugins of Hiplot were visited more than 1,000 times (**Fig. 3C**). The basic heatmap plugin alone has been visited more than 70,000 times, and the correlation heatmap, bubble, boxplot, line-regression, and volcano more than 20,000 times. The UCSCXenaShiny Shiny application in the advanced module was visited 6,307 times at least [13], while the clusterProfiler-based and gene set enrichment analysis (GSEA)-based pathway analyses have been accessed more than 21,000 and 5,000 times respectively. The pdf-collage plugin in the mini-tools module has been visited more than 7,000 times. An increase in user traffic may reflect the potential impact of the web plugins that have been deployed and served in the web service.

**Fig. 3.**
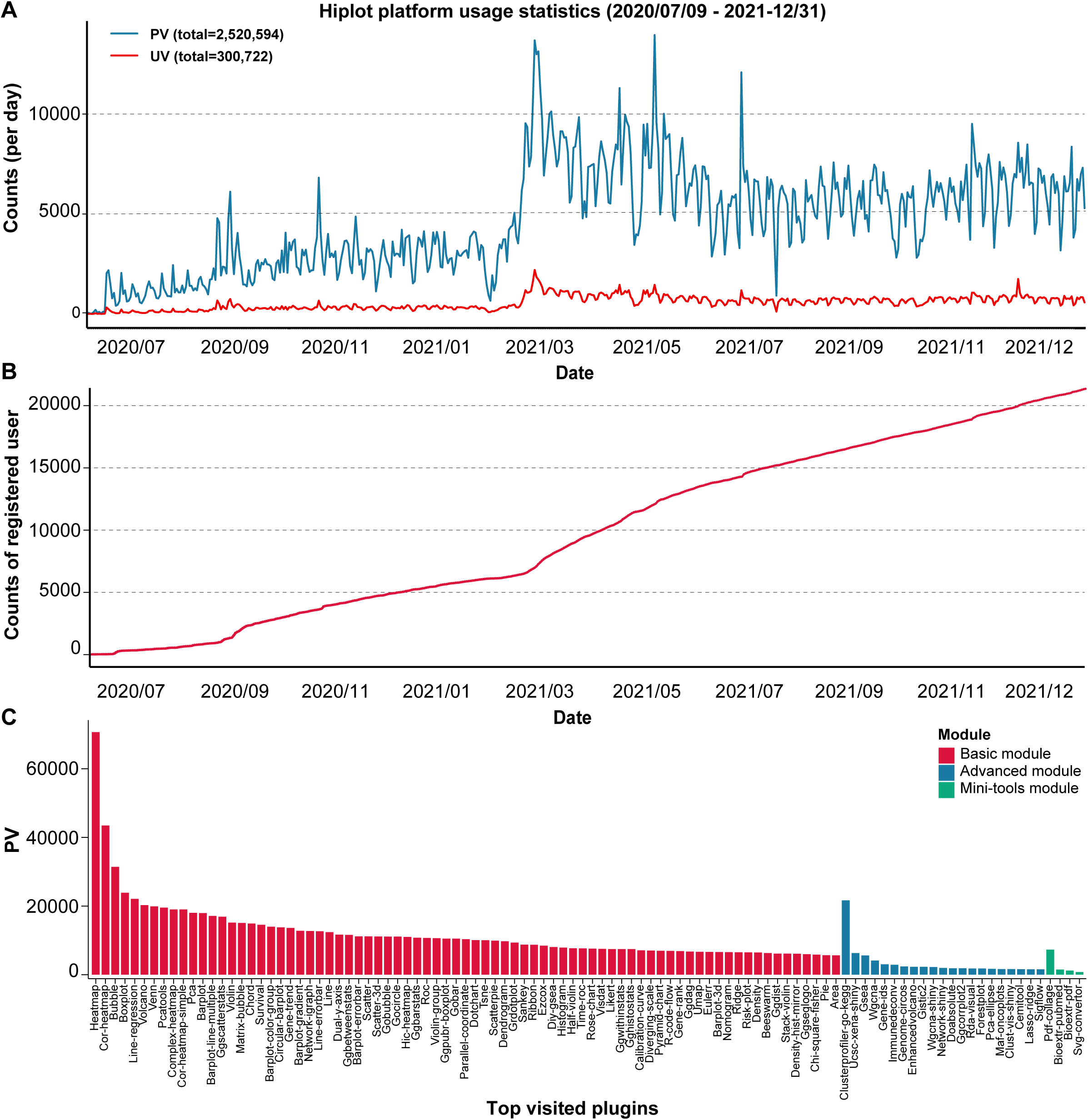
User visit statistics of Hiplot website from July 2020 to December 2021. (A) The line plot shows the PV and UV statistics of Hiplot from July 2020 to December 2021. The website has been visited more than 2,500,000 times, and now reaches 5000 times visits per day. (B) The line plot demonstrates the growth trends of registered users of Hiplot, and more than 22,000 user accounts have been registered. (C) The bar plot indicates the PV of the top-visited plugins in the basic, advanced, and mini-tools module of Hiplot. The order of web plugins is sorted by the module names and PV. The visits of web plugins in the basic module are overall greater than other modules reflecting the common daily demands in the whole community. The heatmap, clusterprofiler-go-kegg, and pdf-collage are the top-visited applications in the basic, advanced, and mini-tools modules respectively. PV, page view. UV, unique visitor.

### Use case 1: basic scientific graphics

One of the common use cases of Hiplot is to draw basic scientific graphics based on tabular data. Users can investigate the correlation of the variables in scatter, chord, line, heatmap plugins, etc. (**Fig. S10A**). At least ten plugins including upset, venn, pie, parliament, waffle, donut, fan, moon charts, tree-map, and flower plot are capable to exhibit the data ratio or topology (**Fig. 1B**), while data distributions can be shown in the histogram, boxplot, violin, ridge, density, area, and bean plots (**Fig. 1B**). Other visualization functions involving data evolution, network relationships, and spatial characteristics also have been developed in the Hiplot, e.g., igraph-based network analysis (**Fig. S10B, C, D**), slopegraph, barplot-line-multiple, and waterfalls plots for viewing the trend of data changes (**Fig. S10E, F**).

Heatmap, the highest high-frequency scientific graphics in the Hiplot website, was chosen to exhibit the common usage of basic visualization tasks (**Fig. 4A**). After entering the plugin page, users can load the demo data via clicking the top/bottom demo button for checking the demo input. In the heatmap plugin, users need to input a numeric data table (one row represents a feature, and one column represents a sample) and optional row/column annotations. The above data tables can be imported from the clipboard, local file, or remote file server. The gene expression matrix of more than 2 megabytes (MB) is recommended to be uploaded in the mode of file path selection with the file uploader. The general parameters of the heatmap plugin can be used to control the width, height, font, theme, color palettes of annotations rows/columns, font size of row/column text, and title. The extra parameters involve the heatmap colors, scale, top variance, the clustering method, the distance measure, and whether display numeric values. Different clustering methods and distance measures could be tried for finding a reasonable result. Default, the heatmap plugin used the Euclidean distance and ward.D2 method in the clustering analysis. If the discrimination of gradient color is not obvious enough, users could try to adjust the extra parameter “scale” for scaling data by row or column. The selection of top variance features is useful for conducting unsupervised hierarchical clustering if the number of inputted features is too large. After submitting the task, the data stream was encoded based on Base64 and then was transferred to the backend services (**Fig. 4B**). Submitted heatmap tasks will commonly be processed by the available plumber workers. Finally, the heatmap output, e.g., JPG and PDF, would be previewed and downloaded in the bottom preview window. To facilitate the next loading of submitted parameters, users can export the parameters as the local JSON file. In addition, the logged-in users can click the sync button in the preview window to permanently save the results to the cloud file manager.

**Fig. 4.**
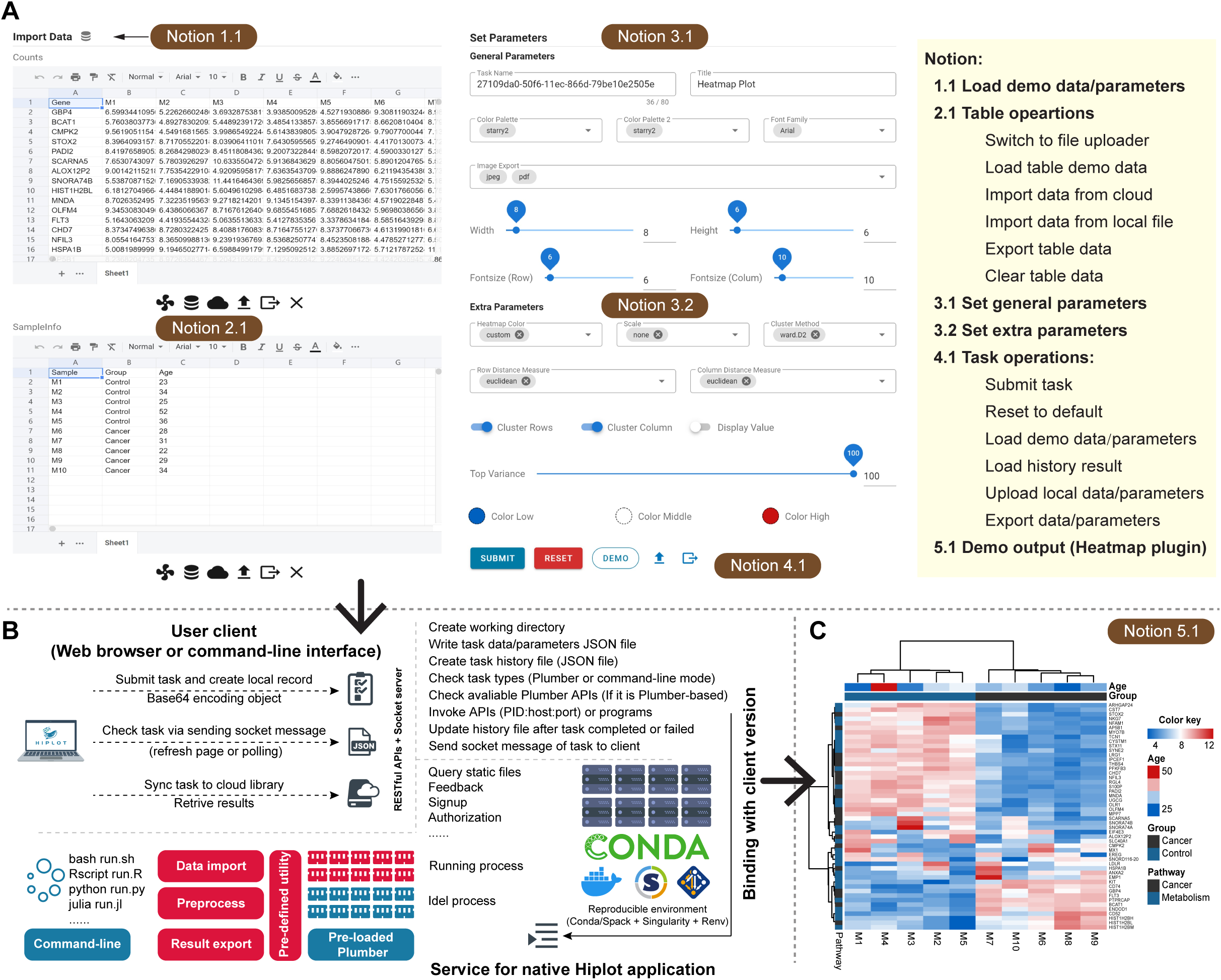
Usage flow and task processing steps through the heatmap plugin. (A) The web interface of the heatmap plugin and the major operation steps for submission of a new task. Tabular data tables are imported and can be previewed and edited in the spreadsheet web component. The general and extra parameters of the heatmap are listed on the right panel. These parameters can control the data pre-processing, clustering steps, and the output style of the heatmap. The parameter “top variance” can be used to select partial genes for conducting the clustering. The parameters of the heatmap are stored in a JSON format data structure and can be directly exported for reproducing the input. The history results in the cloud file manager can be used to reproduce the input and output. (B) The backend response steps and the relevant services processing the submitted task. Submitted data will be encoded using the Base64 algorithm. The available plumber worker processes the heatmap task and the core codes of the heatmap plugin. When the task is finished, the backend returns a history JSON file storing task information and path of output for retrieving the image and logging output. (C) Demo output of heatmap plugin. One column represents a sample and one row represents a gene. Two clusters can be defined in the demo input (cancer vs. control). The row annotations and column annotations respectively show the classification of genes (cancer or metabolism). and sample types (cancer or control).

### Use case 2: omics-based data visualizations in cancer

Dozens of multiple multi-omics data visualizations plugins have been developed in the Hiplot, especially cancer genomics and transcriptomics, which mainly consists of the Shiny applications and the native plugins of Hiplot. In 2019, the open-source project UCSCXenaShiny (**Fig. 5A**) was launched by Openbiox community. In this project, we implement a set of R functions and the Shiny-based web interface allowing users to quickly search, download, explore, analyze, and visualize the dataset from UCSC Xena data hubs [13, 27]. Here, we use the pan-cancer module of UCSCXenaShiny to exhibit that the high expression of *TRH* (thyrotropin releasing hormone) is significantly associated with favorable prognosis and poor prognosis in the acute myeloid leukemia (AML) and glioma patients from The Cancer Genome Atlas (TCGA) database, respectively (**Fig. S11**). In fact, the prognostic significance of any other genes with de-regulated gene expression or sequence mutations in major types of cancer can be explored in this interactive application. Other Shiny-based applications involving cancer omics visualizations could be found via clicking the “Shiny” and other related tags in the advanced module of the Hiplot website, such as genome-wide association study (GWAS)-related Shiny plugins (**Fig. S12, S13**).

**Fig. 5.**
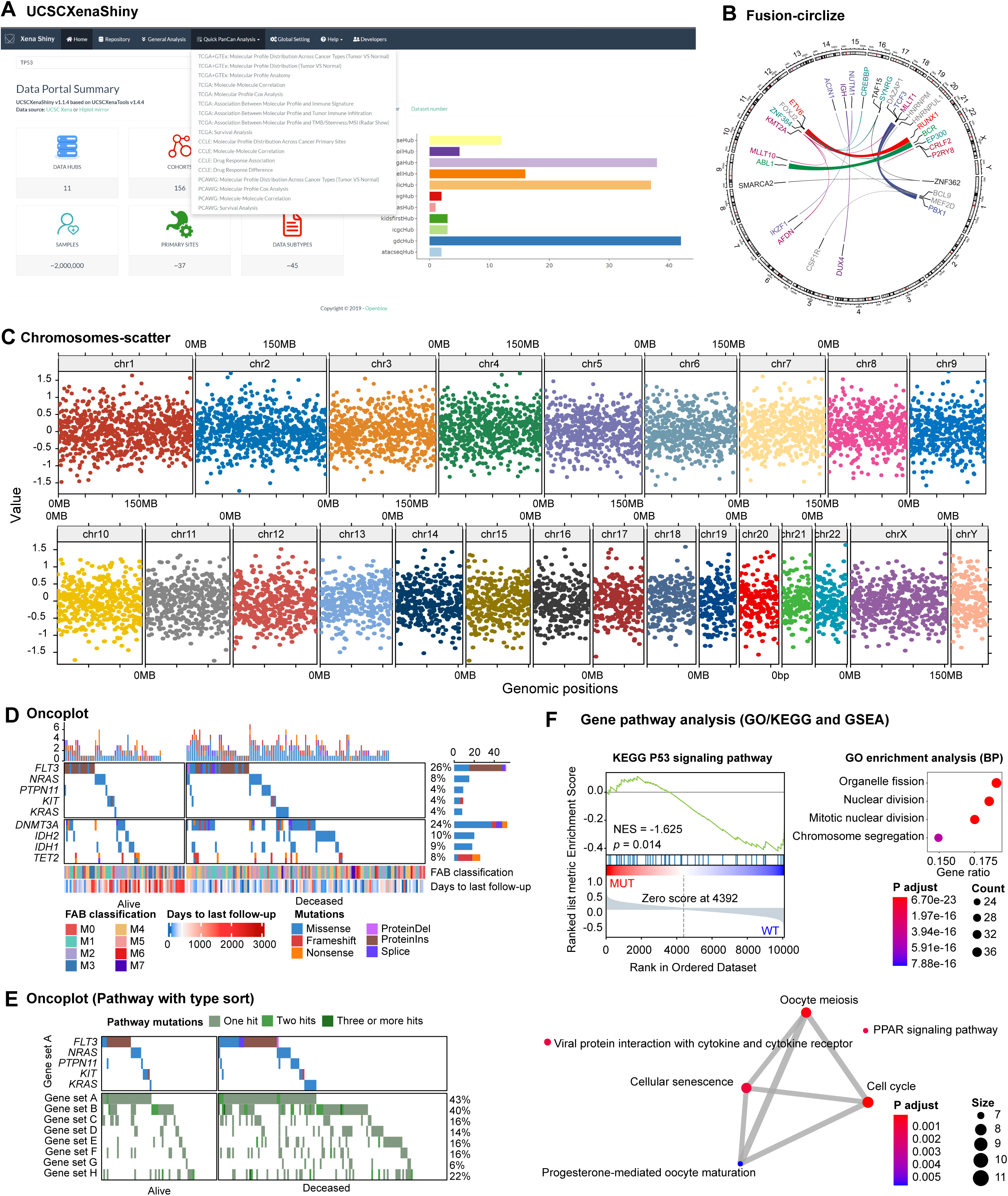
Representative use cases of omics-based visualization functions in Hiplot. (A) Screenshot of UCSCXenaShiny application, which is an R shiny-based application for interactively mining the published datasets from UCSC Xena Hub. (B) Demo output of the fusion-circlize plugin. It shows a part of known gene fusions including *ETV6*-*RUNX1, BCR*-*ABL1, DUX4* fusions, *ZNF384* fusions, *MEF2D* fusions, and *KMT2A* fusions in B-cell acute precursor lymphoblastic leukemia (BCP-ALL). Different chromosomes are ordered clockwise. Different chromosomes are ordered clockwise. The gene fusions are linked between chromosomes using ribbons. The color and the width of the link/text indicate the classes and frequency of the specific fusion genes. (C) Demo output of chromosomes-scatter plugin, which shows the simulated numeric values in different chromosomes with colors (D) Oncoplot shows selected mutant genes in the acute myoid leukemia (AML) cohort from The Cancer Genome Atlas (TCGA) database. The patients are split into two parts according to their survival status. Different mutation types are labeled in different colors. The bottom box draws the meta-annotation of patients. (E) Another Oncoplot with extra the gene pathway rows and the patients are sorted by mutation types. (F) The demo outputs of gsea and clusterprofile-go-kegg plugins. The NES and p-value are shown in the GSEA output, which is regenerated to the PDF version. The bubble plot and network plot show the enriched GO pathways based on the demo data of clusterprofile-go-kegg plugin. KEGG, Kyoto Encyclopedia of Genes and Genomes. UCSC, University of California, Santa Cruz. TCGA, The Cancer Genome Atlas. GO, gene ontology. GSEA, gene sets enrichment analysis. NES, normalized enrichment score.

Apart from Shiny-based applications, several JSON-based native plugins of Hiplot can be used to interactively visualize large-scale cancer omics data. The fusion-circlize plugin displays the demo gene fusions in B-cell precursor acute lymphocytic leukemia (BCP-ALL) at the chromosomal level [28], such as *BCR*-*ABL1, ETV6*-*RUNX1, DUX4* fusions, *ZNF384* fusions, *MEF2D* fusions, *KMT2A* fusions, and *NUTM1* fusions (**Fig. 5B**). In this plugin, the colors and ribbon width of fusion genes can be customized. The chromosomes-scatter plugin visualizes the numeric value using scatters of chromosomes (**Fig. 5C**), which may help to display the level of gene copy numbers and gene expressions on a large genome-scale. The oncoplot plugin was developed based on the ComplexHeatmap [2], which was used to display the selected mutant genes and gene pathways of patients from the TCGA LAML cohort (**Fig. 5D, E**). Based on the published variants data of patients with BCP-ALL [28], we invoked multiple plugins of Hiplot and validated that the major fusion genes and chromosomal abnormalities are mostly mutually exclusive in BCP-ALL, while *CRLF2* fusions, *DUX4* fusions, and *BCR*-*ABL1* significantly coexist with the *JAK2, MYC*, and *RUNX1* sequence variants respectively (**Fig. S14**). The coexistence mutual exclusion analysis is suggested to be done in the discover-mut-test plugin with better performance in recognition of exclusive events compared with the classical Fisher’s exact test [23].

The transcriptomic data can be visualized in several basic graphics, such as heatmap, boxplot, volcano, and pseudo-enhanced-ma, for displaying the gene expression level, characteristic genes, and conducting unsupervised clustering (**Fig. 1B**). For example, in the volcano plugin, users can interactively add the selected gene labels of differentially expressed genes (DEGs). The demo of consensus clustering based on gene expression data was completed in the cola plugin (**Fig. S15**). It provides the interface to get stable clusters via data sampling and integrating different top-value and clustering methods [20]. In addition, we used the complex-heatmap plugin to correlate multi-omics features using gene expression, gene mutations, and clinical features (**Fig. S16**) [2].

In the clusterprofiler-go-kegg plugin, we performed a multi-group pathway enrichment analysis of DEGs in a single run based on the demo multi-columns data (**Fig. 5F**). It supports inputting the gene symbol, ensemble id, and gene id format data and will merge the multiple graphics and table output in a single PDF and Excel file. The gsea plugin is a complete web implementation of the original Broad GSEA command-line program. Compared with the desktop version, this web plugin allows users to simultaneously compare multiple subgroups and multiple gene sets (**Fig. 5G**). Besides, we constructed the first web interface of immunedeconv R package, which allows users to perform the TME analysis and to calculate the immune cell fraction based on multiple algorithms including quantiseq, TIMER, CIBERSORT, MCPCounter, xCell, and EPIC (**Fig. S17**) [29-35].

### Use case 3: dimensional reductions and clinical data visualizations

Dimension reduction analysis (DRA), such as the principal component analysis (PCA), t-distributed stochastic neighbor embedding (tSNE), and uniform manifold approximation and projection (UMAP), etc., selects the most important dimensions from multidimensional data according to the sorting method of features or distances as representative data for subsequent analysis, thus helping to identify potential new types of cells/patients with biological/clinical significance. Here, we used multiple DRA methods to visualize the well-known iris dataset and showed the known classes in different data spaces after dimensionality reduction (**Fig. S18**). In the clinical data visualizations tasks, we used the ezcox plugin to conduct a batch of Cox modeling using a multivariate survival risk model with control variable [36] (**Fig. 6A, B**). The metawho plugin is a simple web implementation of the “Meta-analytical method to Identify Who Benefits Most from Treatments” for conducting the meta-analysis tasks (**Fig. 6C**) [37]. Meanwhile, the risk-plot (**Fig. 6D**) and survival (**Fig. 6E**) plugins were used to visualize the disease risk models and survival data of patients. The risk-plot plugin can display the correlation between survival status and risk factors. The patients were sorted by the risk scores from low to high. In the nomogram plugin, we established the demo predictive risk score based on the lung dataset in the survival package (**Fig. 6F**).

**Fig. 6.**
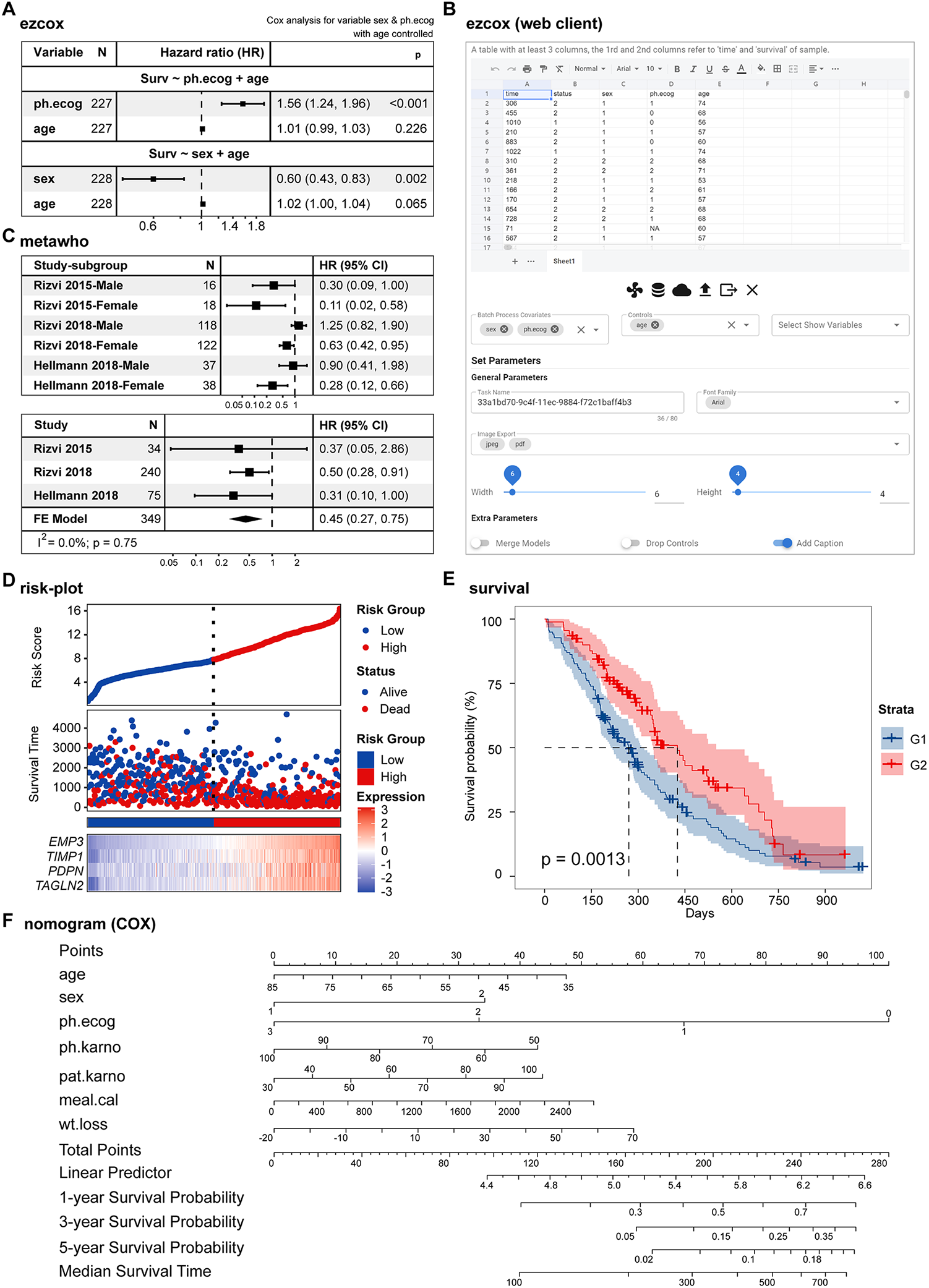
Representative use cases of clinical data visualization in Hiplot. (A) and (B) respectively shows the demo forest output using the lung dataset in the survival package and the web interface of the ezcox plugin. The COX models show that the sex and ph.ecog variables are respectively associated with high risk (HR:1.56, CI: 1.24-1.96) and low risk (HR:0.60, CI: 0.43-0.83) in the lung when age is used as the control variable. (C) Demo output of the metawho plugin. The HR of study subgroups and studies are displayed in the table. (D) Demo output of the risk-plot plugin. One point represents one patient, which is divided into high risk (red) and low risk (blue) subgroups according to the median value of risk scores. Dead patients are shown in red points in the middlebox. The expression of feature genes including *EMP3, TIMP1, PDPN*, and *TAGLN2*, displayed in the bottom box. (E) Demo output of the survival plugin. It indicates the three years survival curve in G1 (blue) and G2 (red) subgroups of simulated data. (F) Nomogram plot that can be used for predicting the prognosis in the lung dataset of the survival package. The p-value is calculated based on the log-rank test. HR, hazard ratio. CI, confidence interval.

## Conclusion

In summary, a comprehensive and easy-to-use cloud service, Hiplot, is proposed in this work for interactively conducting biomedical data visualization. The concise user interfaces and efficient interaction methods of Hiplot have minimized the learning and usage costs of lightweight visualization tasks for users without programming skills. The demo and real datasets show the web-based visualization functions of Hiplot. Our work provides an important and useful online resource for biomedical researchers and other data scientists.

## Availability and requirements

**Project name:** Hiplot

**Project home page:** https://hiplot.com.cn

**Any restrictions to use by non-academics:** license required

## List of abbreviations

CLI: command-line interface
JSON: JavaScript Object Notation
RESTful: resource representational state transfer
APIs: application programming interfaces
GO: gene ontology
KEGG: Kyoto Encyclopedia of Genes and Genomes
UCSC: University of California, Santa Cruz
TCGA: The Cancer Genome Atlas
GSEA: gene sets enrichment analysis
GWAS: genome-wide association study
HR: hazard ratio
CI: confidence interval
DEGs: differentially expressed genes
TME: tumor microenvironment
DRA: dimensional reduction algorithm
BCP-ALL: B-cell precursor acute lymphocytic leukemia
PCA: principal component analysis
tSNE: t-distributed stochastic neighbor embedding
UMAP: uniform manifold approximation and projection

## Declarations

### Ethics approval and consent to participate

Not applicable.

### Consent for publication

Not applicable.

### Availability of data and materials

The website can be freely accessed via http://hiplot.com.cn. The available open-source code of the website are located at http://github.com/hiplot.

### Authors’ contributions

L.J.F, W.M.J, C.S.J, C.Z, S.Y, W.R.X, L.X.S designed and supervised the study. L.J.F, W.M.J, M.B.B, and W.S.X designed and implemented the native Hiplot framework and infrastructure of the web service. L.J.F and M.B.B contributed the command-line tool HCTL. L.J.F, W.M.J, M.B.B, W.S.X, D.W, and X.H.S developed the major applications of Hiplot. D.S.Q, L.J.C, and B.Z.W developed the minor applications of Hiplot. L.J.F, M.B.B, and W.S.X wrote the manuscript and other co-authors critically reviewed and modified the manuscript.

## Acknowledgements

The authors are grateful and indebted to the members of the Galaxy, UCSC Xena, UCSCXenaShiny project, Openbiox community, Shanghai Tengyun Biological Technology Co., Ltd., and other developers of open-source scientific applications. Thanks are due to Yang Xu of the National Earth System Science Data Center of China for providing help in developing geoinformatics applications. Useful suggestions given by Yutong Ding and other members of the China National GeneBank are also acknowledged.

