## Supplementary Figures for "Hiplot: a comprehensive and easy-to-use web service boosting publication-ready biomedical data visualization"

**Affiliations of authors:**

[1] Shanghai Institute of Hematology, State Key Laboratory of Medical Genomics, National Research Center for Translational Medicine at Shanghai, Ruijin Hospital Affiliated to Shanghai Jiao Tong University School of Medicine, Shanghai, China; [2] School of Life Sciences and Biotechnology, Shanghai Jiao Tong University, Shanghai, China; [3] College of Fisheries, Guangdong Ocean University, Zhanjiang, Guangdong, China; [4] School of Life Science and Technology, ShanghaiTech University, Shanghai, China; [5] Zhongshan School of Medicine, Sun Yat-sen University, Guangzhou, China; [6] Shanghai General Hospital, Shanghai Jiao Tong University School of Medicine, Shanghai, China; [7] Department of Histology, Embryology, Genetics and Developmental Biology, Shanghai Key Laboratory for Reproductive Medicine, Shanghai Jiao Tong University School of Medicine, Shanghai, China; [8] Department of Urology, First Hospital of Shanxi Medical University, Taiyuan, China; [9] College of Food Science, Sichuan Agricultural University, Yaan, China; [10] Department of Neurosurgery, The Second Affiliated Hospital of Dalian Medical University, Dalian, China; [11] State Key

Laboratory for Diagnosis and Treatment of Infectious Diseases, Collaborative Innovation Center for Diagnosis and Treatment of Infectious Diseases, The First Affiliated Hospital, Zhejiang University School of Medicine, Hangzhou, China; [12] Department of Endocrinology and Metabolism, Zhongshan Hospital, Fudan Institute for Metabolic Diseases, and Human Phenome Institute, Fudan University, Shanghai, China; [13] Qinghai Academy of Animal and Veterinary Science, State Key Laboratory of Plateau Ecology and Agriculture in the Three River Head Waters Region, Qinghai Provincial Key Laboratory of Adaptive Management on Alpine Grassland Qinghai University, Xining, China;[14] National Key Laboratory of Plant Molecular Genetics, CAS Center for Excellence in Molecular Plant Sciences, Shanghai Institute of Plant Physiology and Ecology, Shanghai Institutes for Biological Sciences, Chinese Academy of Sciences, Shanghai, China; [15] University of the Chinese Academy of Sciences, Beijing, China; [16] Faculty of Health Sciences, University of Macau, Taipa, Macau, China; [17] Department of Gastroenterology, Ruijin Hospital, Shanghai Jiao Tong University School of Medicine, Shanghai, China; [18] These authors contributed equally: Jianfeng Li, Benben Miao, Shixiang Wang, Wei Dong, Houshi Xu, Chenchen Si;

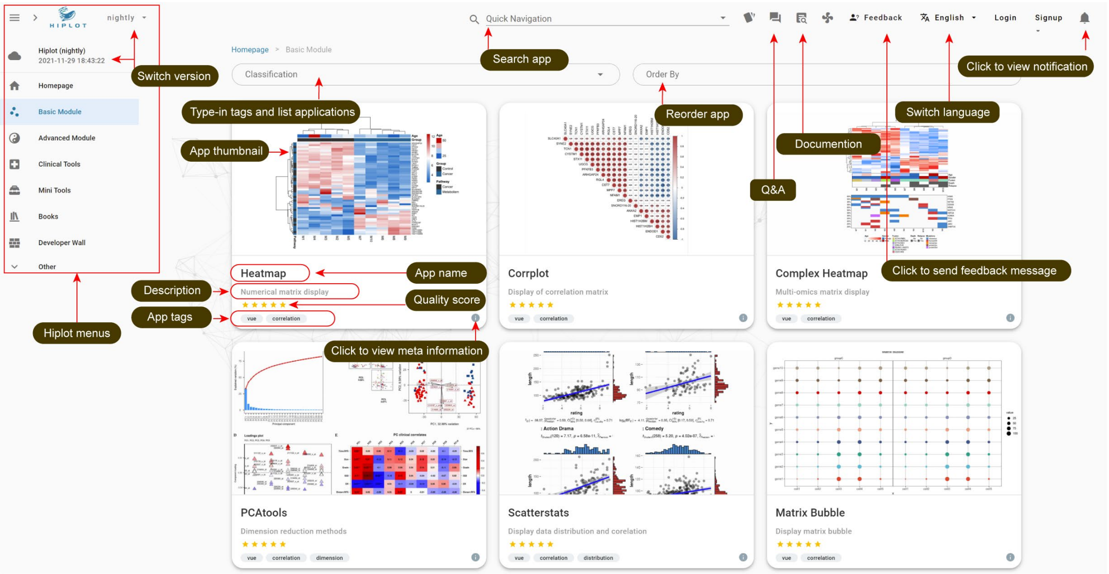


**Fig. S1. Layout and functional buttons of Hiplot website in the card list view of modules.** Left menu can be used to switch the modules of Hiplot. Right side of page list the card list of plugins. The basic meta information including the name, short descriptions, tags, and quality score are shown in the cards. The plugins can be filtered by "Classification" text field and sorted by "Order By" text field on the top of plugin panel.

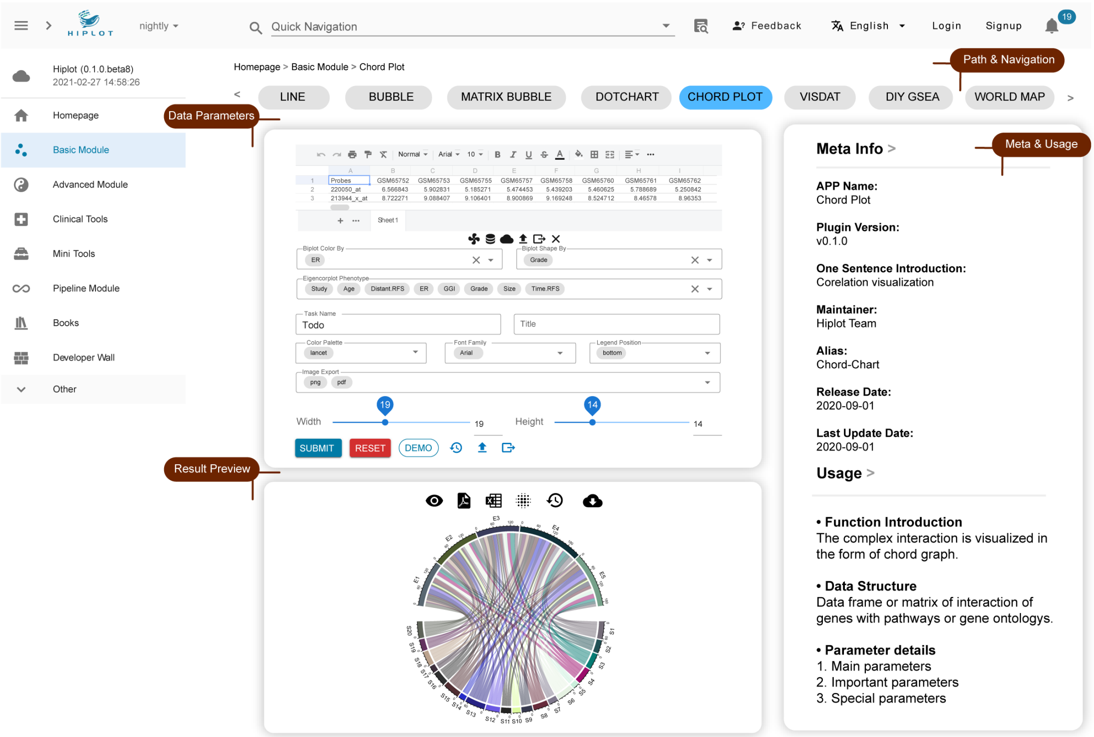


**Fig. S2. Layout overview of a simple Hiplot plugin.** Three windows including data/parameters, preview, and documentation are provided in the working page of the website. The applications can be switched using the top path or via the search function in the “Quick Navigation” field. The task records window and its menu are not shown in this schematic diagram, which can be switched in the preview window.

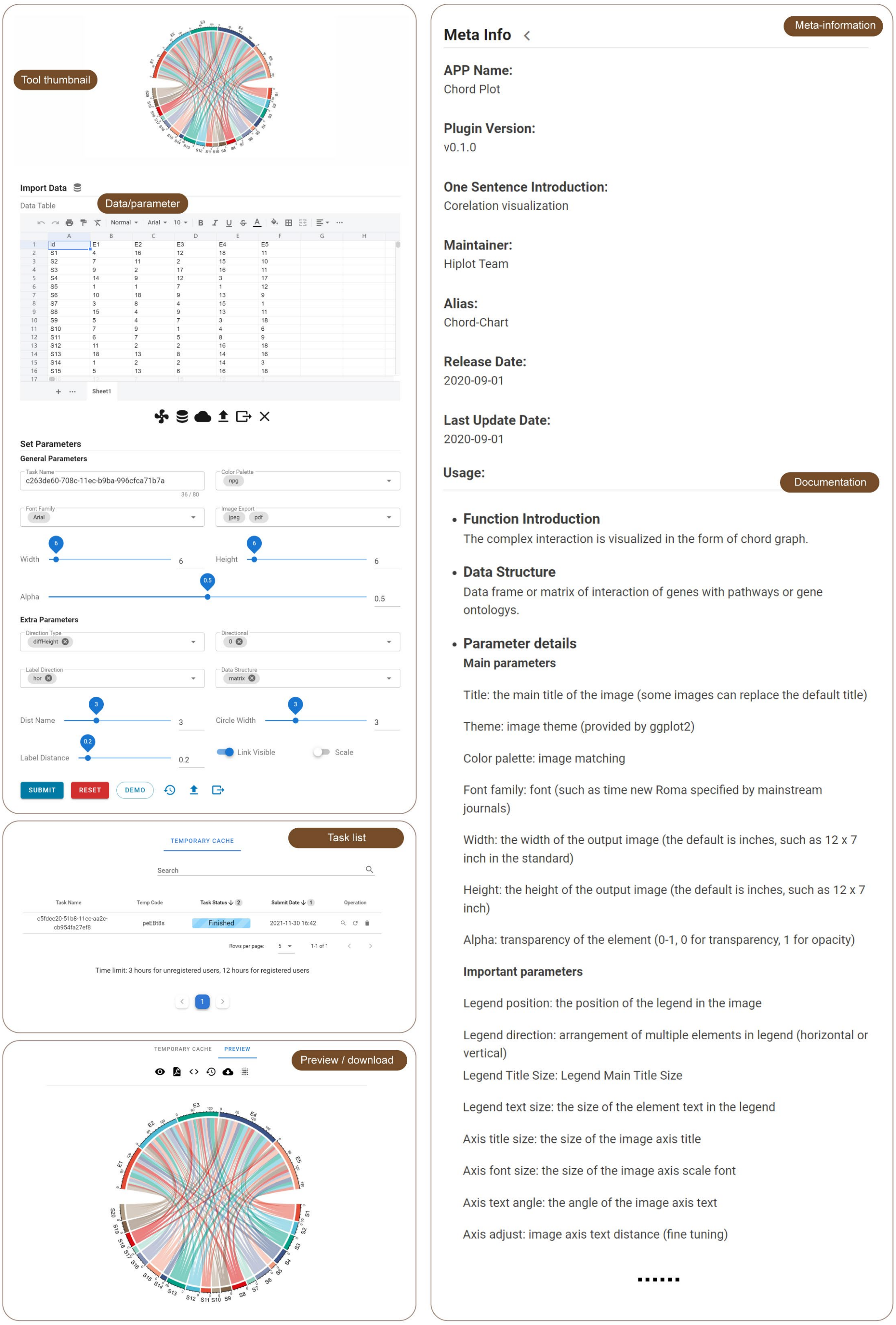


**Fig. S3. Layout and functional area of the Hiplot plugins.** The long screenshot indicates the layout of chord plot plugin of the basic module. The layout includes the tool thumbnail, data/parameter, task list, preview/download, meta-information, and documentation. The content of task list and preview window are shown separately. When a new task is submitted, a new line would be added to the task list to show the task status.

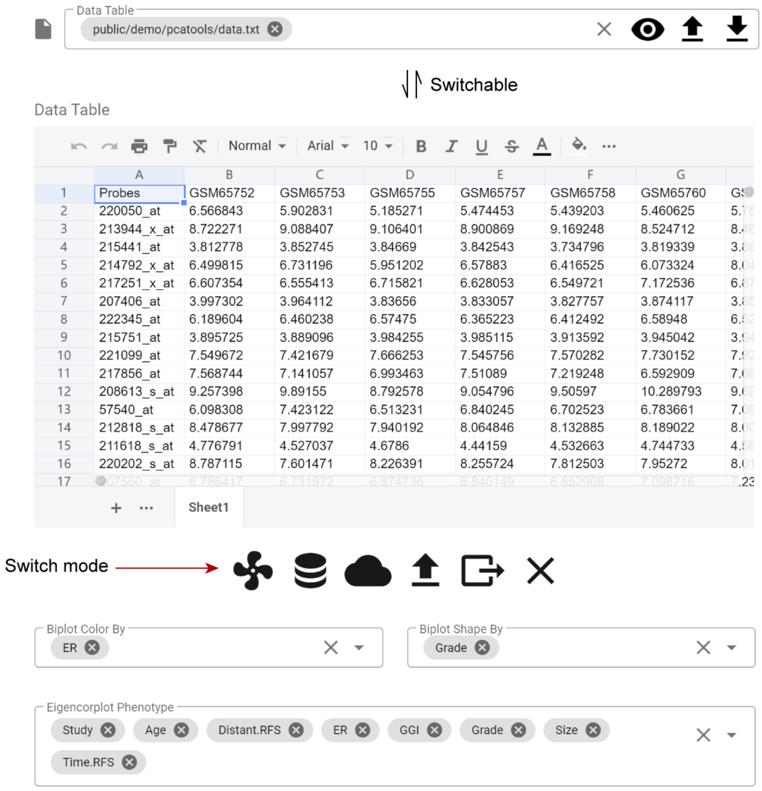


**Fig. S4. Switchable data importing from file path selection and the online table editor.** The input field on the top is the file path selection mode, which can be used to upload large-size files. The spreadsheet on the bottom is the tabular mode for loading small-size data. These two modes can switch to each other via clicking the first button on the table bottom when users have logged in the website. The columns selection fields provide users an easy way to transfer custom commands to backend operations.

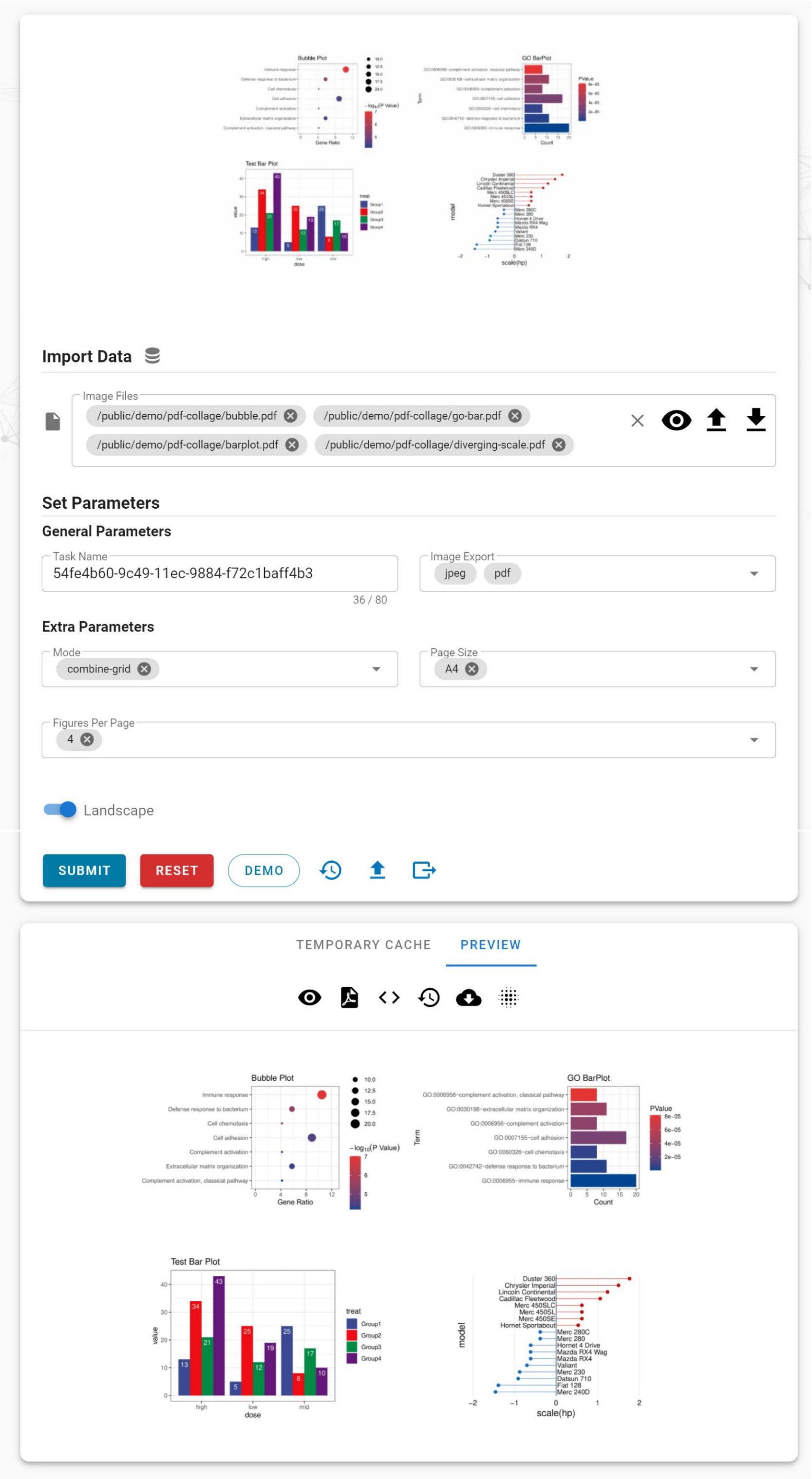


**Fig. S5. Web interface and demo output of pdf-collage plugin.** Before using this plugin, users need to upload multiple figures from local or select the history results from cloud library. The items per page, e.g., 4 per page, the page orientation can be modified to fit user's demands.

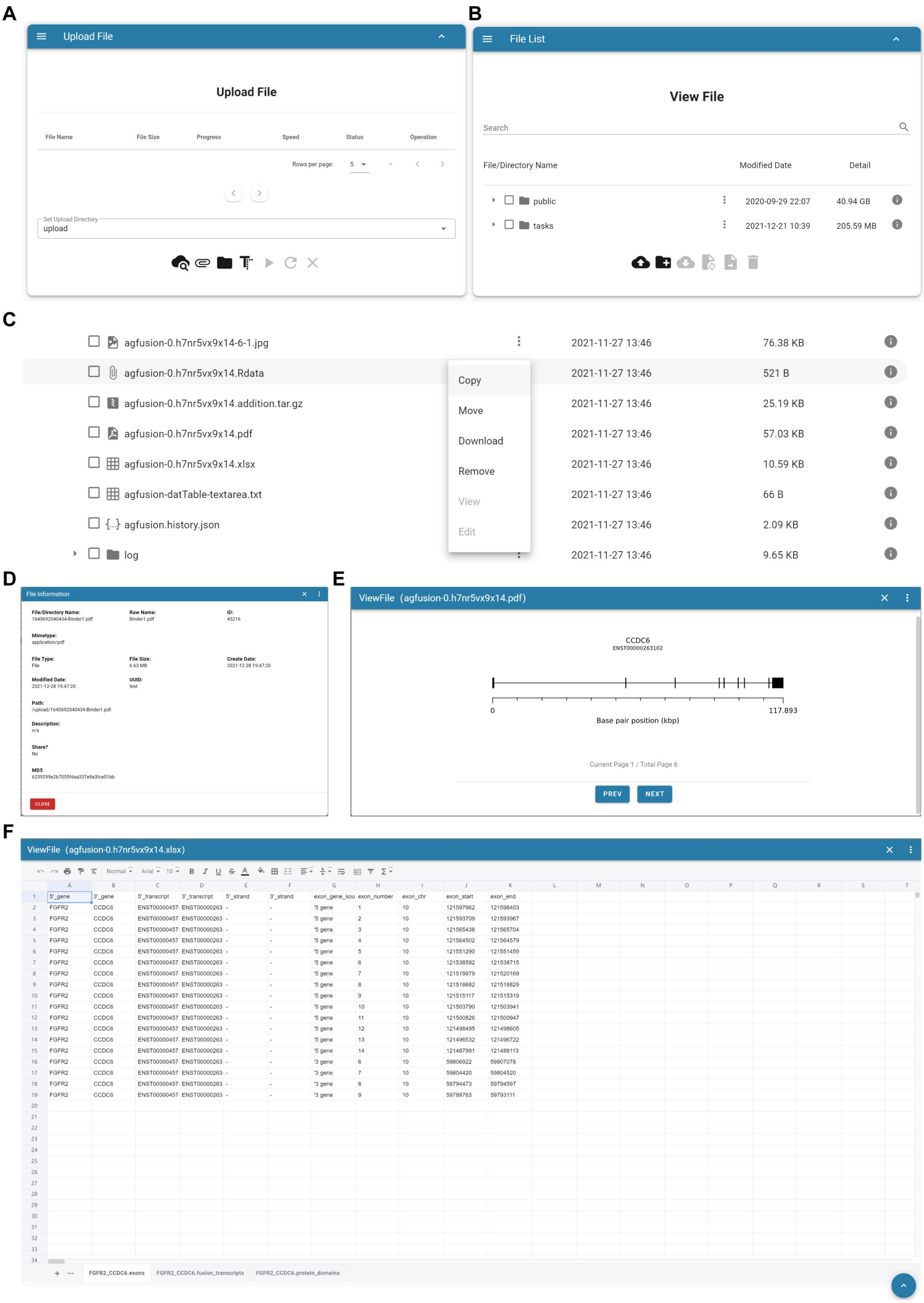


**Fig. S6. File manager module of Hiplot.** Screenshots show (A) the window for file upload, (B) the tree view of uploaded file list, (C) available file operations including copy, move, download, remove, view, and edit, (D) the fields and meta-information of uploaded files, (E) preview window for image, and (F) preview window with a spreadsheet for table data.

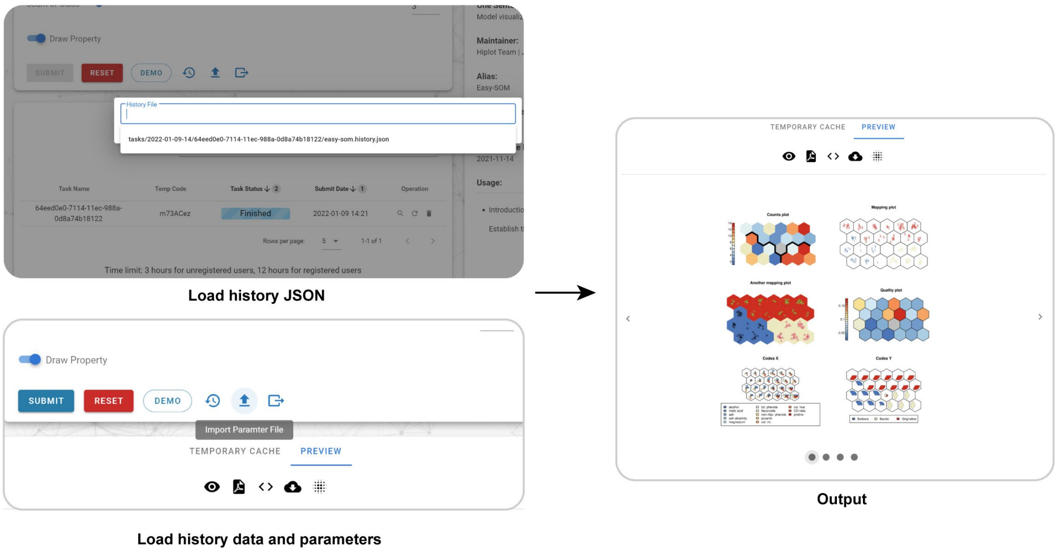


**Fig. S7. Two optional methods for reproducing the history input and output.** The screenshot shows how the demo outputs of the easy-som plugin are reproduced based on the local *Data JSON* object or remote history file.

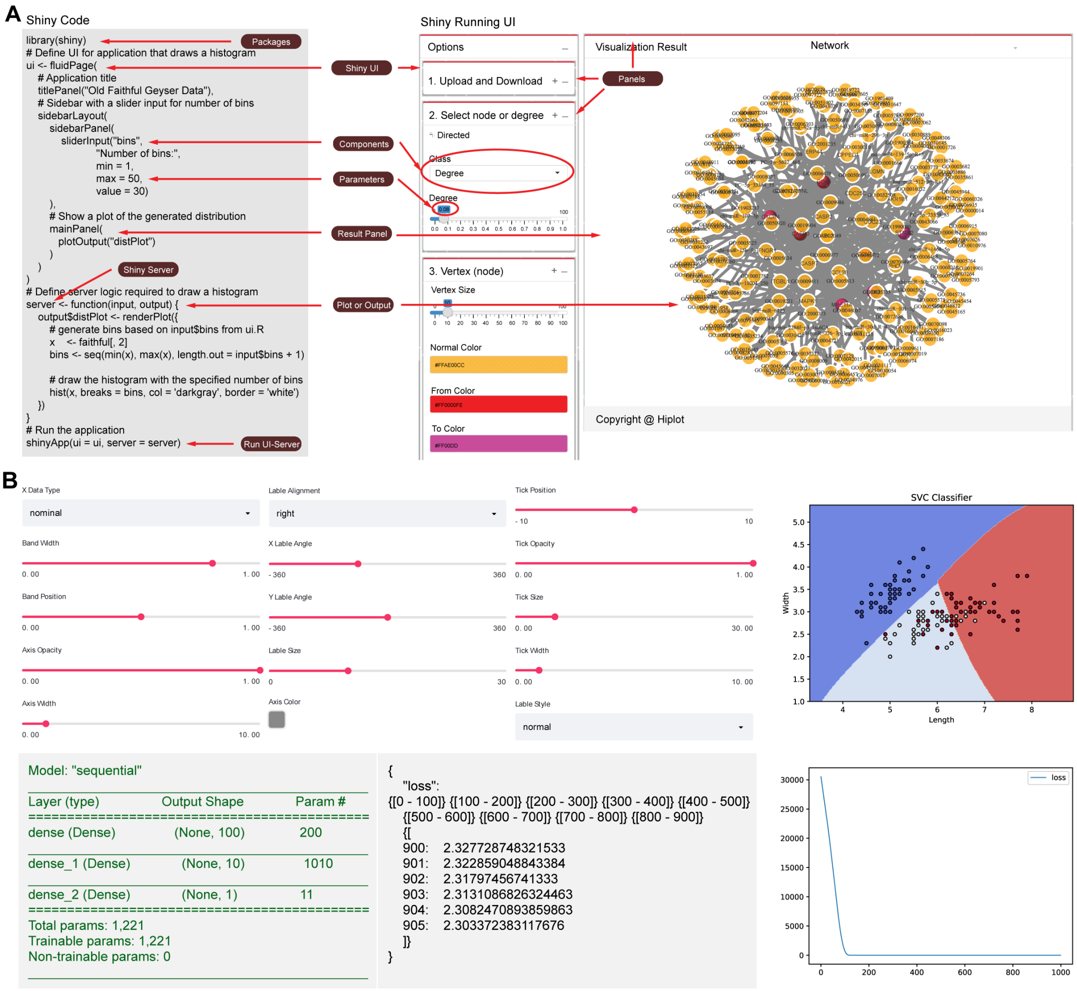


**Fig. S8. UI Layout based on the R Shiny and Python Stremllite framework.** (A) The R Shiny-based network-shiny plugin. Left panel shows the code snapshot and right panel displays the front-end interface and demo output. (B) The Python Stremllite-based strml-svm-classifier plugin. Top-left panel shows the front-end interface. The bottom-left prints the logging of training process and the right shows the visualized outputs.

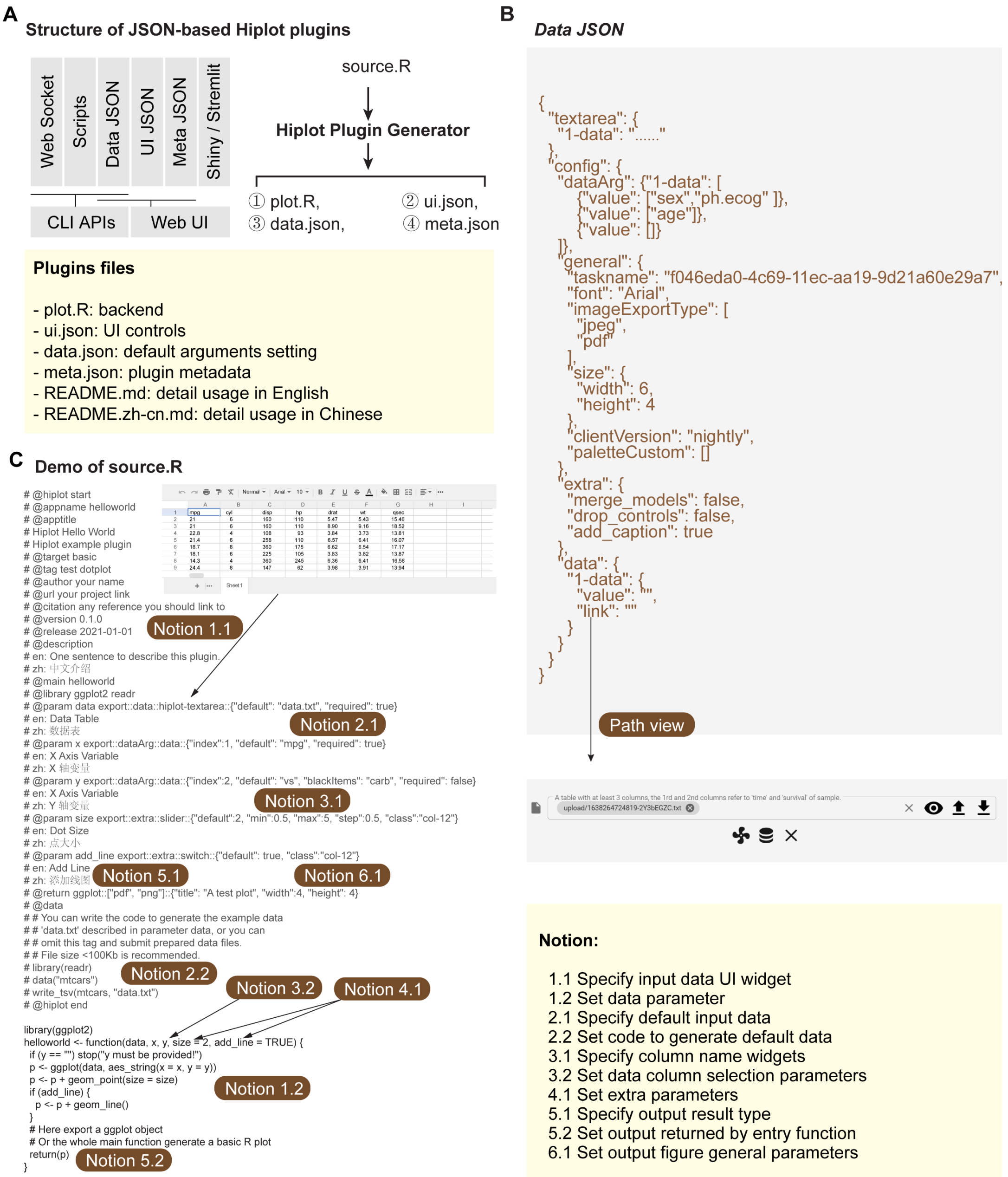


**Fig. S9. JSON-based plugin components and its generator.** (A) The standard file structure of the JSON-based Hiplot plugin includes the plot.R script, ui.json, data.json, meta.json, and documentation Markdown files. The JSON files and plot.R can be automatically rendered from the source.R contains the fields storing the required information (B) Code snapshot of *Data JSON* shows the data fields and demo values of the JSON-based plugin. It can be exported from the plugins page. (C) Snapshot of demo code for generating the Hiplot plugin based on the HiSub program.

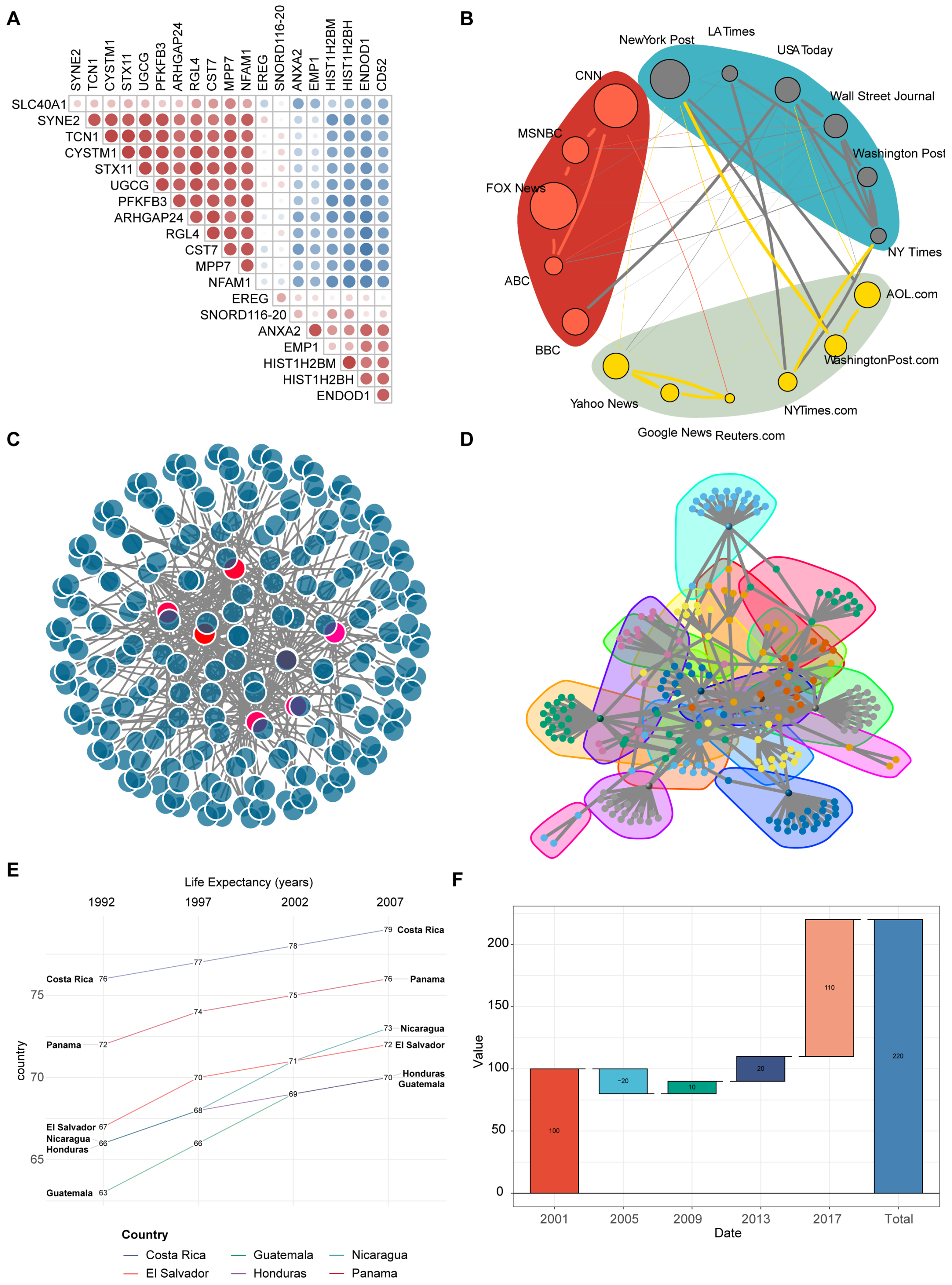


**Fig. S10. Use cases of basic graphics: network and evolution analysis.** Demo outputs of (A) corrplot, (B) network-igraph, (C) network-shiny, (D) community-shiny, (E) slopegraph, and waterfalls plugins. In the corrplot, red represent positive correlation and blue represent negative correlation, while the size of circle indicates the degree of correlations. The network visualizations are constructed based on the igraph R package. The slopegraph and waterfalls shows the values in different time points.

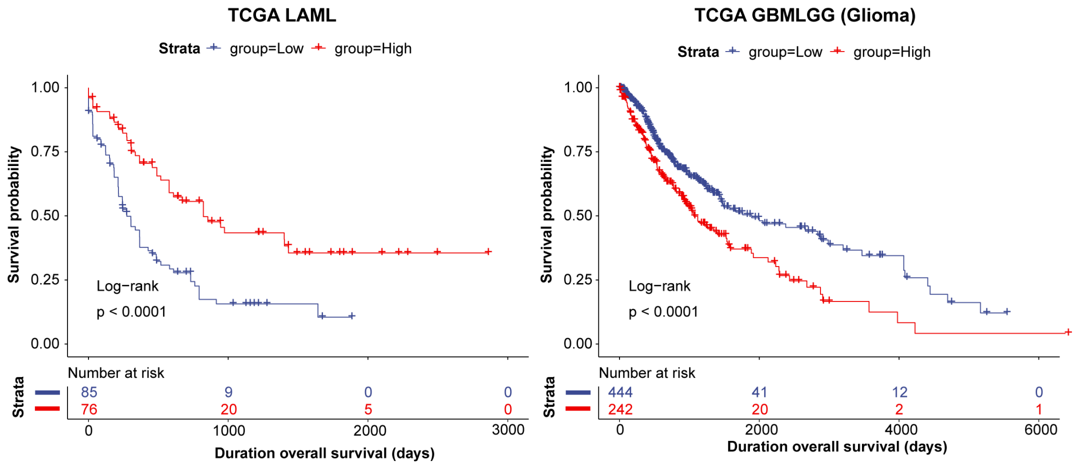


**Fig. S11. Use cases of omics-based data visualizations: the prognosis significance of high gene expression of TRH in public TCGA LAML and GBMLGG cohort.** The left and right panel show the long-term survival rates in the patients from the TCGA LAML and GBMLGG cohorts, respectively. Red lines and blue lines represent the subgroups with high and low expression of *TRH*. TCGA, The Cancer Genome Atlas. GBMLGG, glioma.

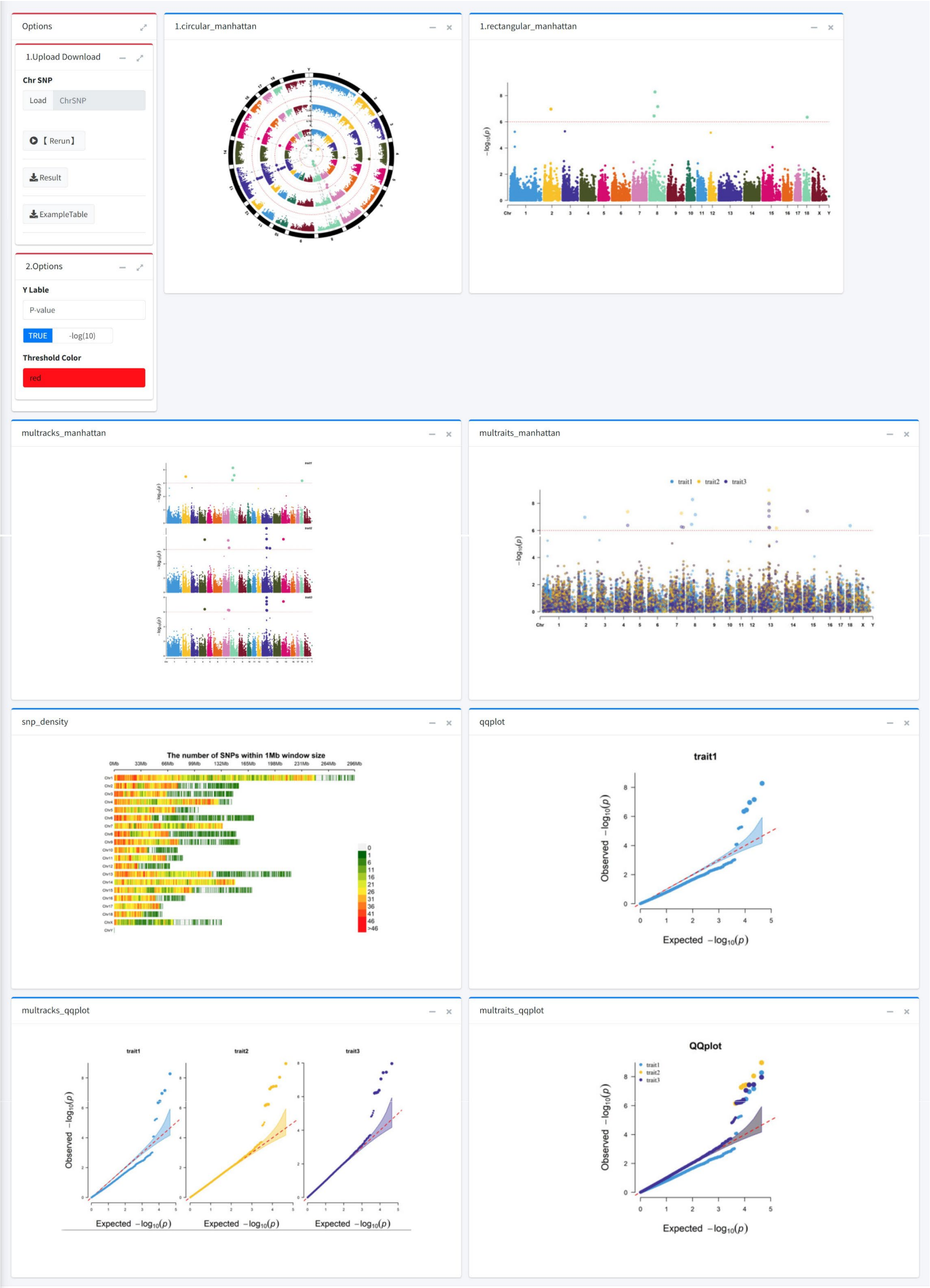


**Fig. S12. Use cases of omics-based data visualizations: the R Shiny-based cmplot application.** Based on the advantages of CMplot R package in analyzing Genome-Wide Association Study (GWAS) data, we further use Shiny framework to develop a user-friendly interface for it, and give interfaces such as program data upload, parameter modification, function automation, image (PDF, SVG, JPEG, PNG, TIFF) download, etc., so that users can calculate online at any time. The data structure is a standard SNP matrix, which include respectively (SNP ID, Chromosome, Position, Trait1, Trait2, Trait3 ...) columns. The program allows users to run circular-Manhattan, single-track Manhattan, multi-tracks Manhattan (all traits in separated axes, all traits in one axes) , SNP density visualization, single-track Q-Q, and multi-tracks Q-Q (all traits in separated axes, all traits in a axes) corresponds to the above six results respectively.

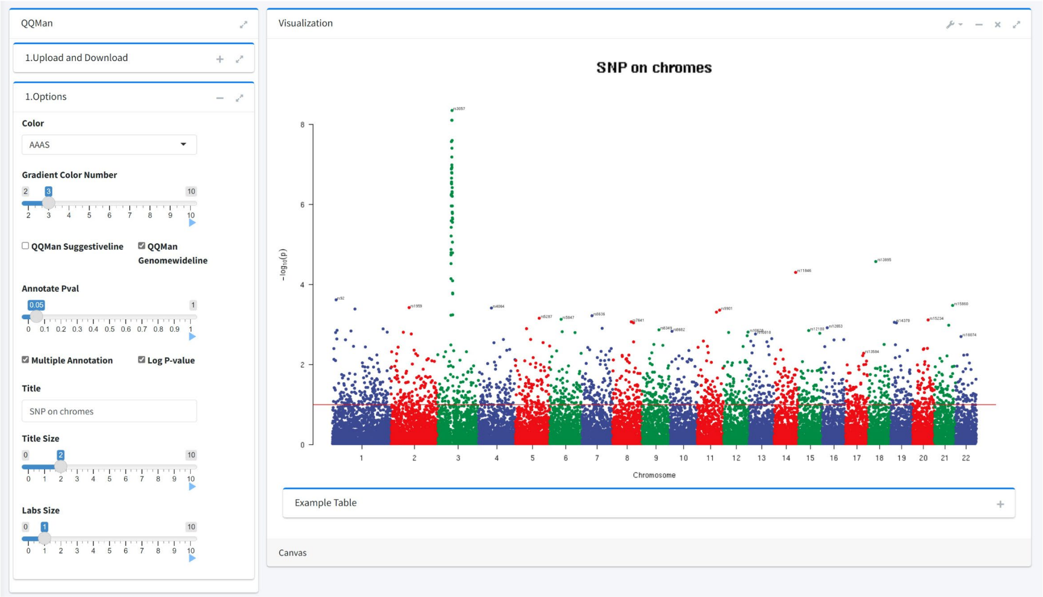


**Fig. S13. Use cases of omics-based data visualizations: the R Shiny-based Manhattan application.** In order to meet the needs of standard and more complex SNP screening and visual analysis, we integrate the QQMan and Manhattan R packages, and combine ggsci package for their visual color matching. The developed interface program divides the complex parameters of functions into default and user-defined according to the analysis needs, allowing users to analyze data more freely and at low cost. The data structure is a standard SNP matrix, which include respectively (SNP ID, Chromosome, Position, Trait1, Trait2, Trait3 ...) columns.

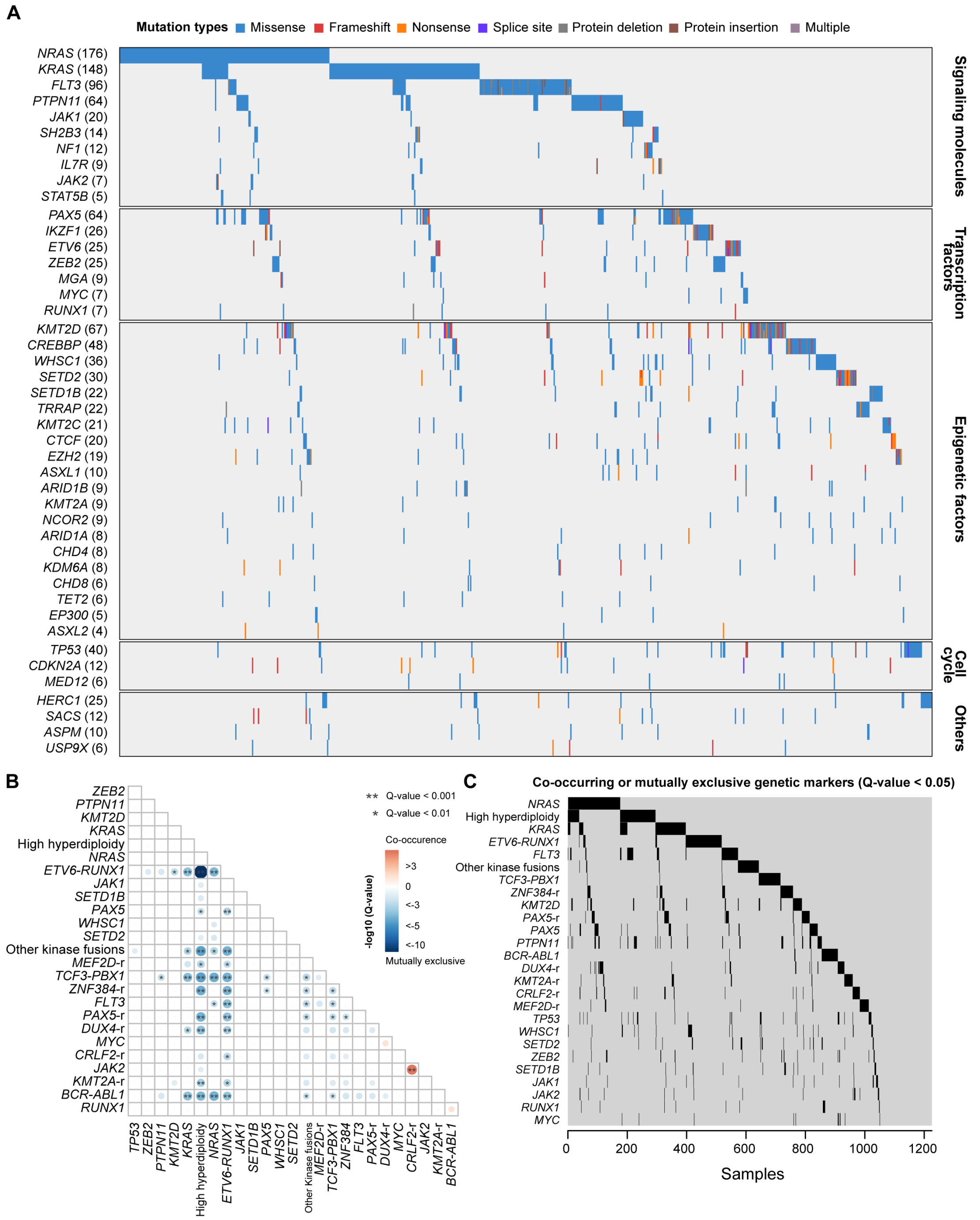


**Fig. S14. Use cases of omics-based data visualizations: mutation landscape and the coexisting and mutually exclusive events of 1,223 patients with BCP-ALL.** (A) Oncoplot shows the mutant genes with sequence variants in patients with BCP-ALL. Different mutation types are filled by different colors. The patients are sorted, and the genes are classified according to gene pathways. (B) Heatmap demonstrates potential coexisting (red) and mutually exclusive events (blue) pairs of gene fusions and sequence variants in BCP-ALL, which was calculated in the discover-mut-test plugin.

(C) Oncoplot based on the paired mutant genes or gene fusions with at least one coexisting or mutually exclusive event.

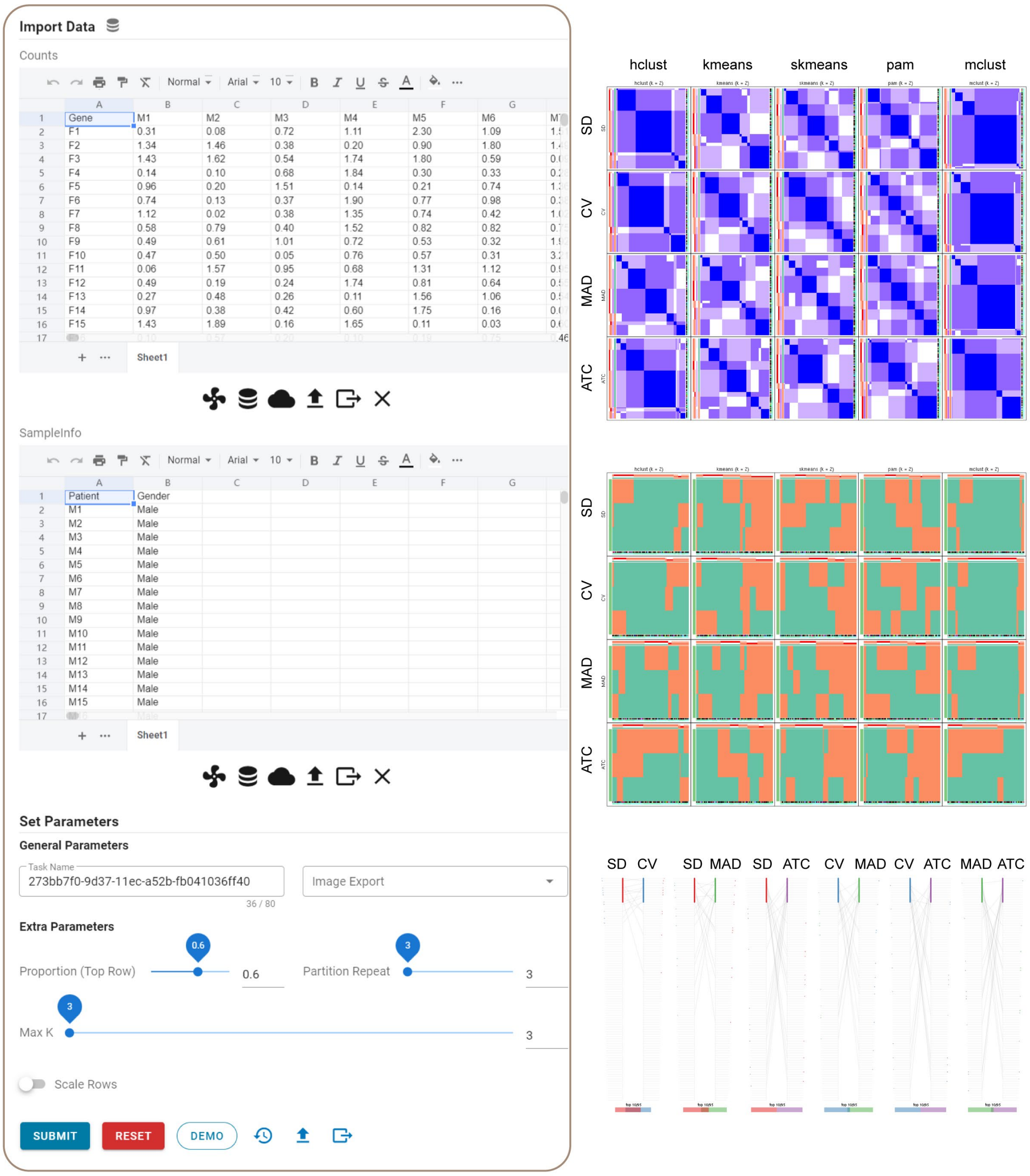


**Fig. S15. Use cases of omics-based data visualizations: consensus clustering based on the cola plugin.** Left panel shows the web interface of cola plugin. Two data table are required including the gene expression data and the annotation table of samples. Right panel shows the partial outputs including consensus heatmap, membership heatmap, and the top rows overlap in cola consensus clustering based on different settings for top-value and clustering methods.

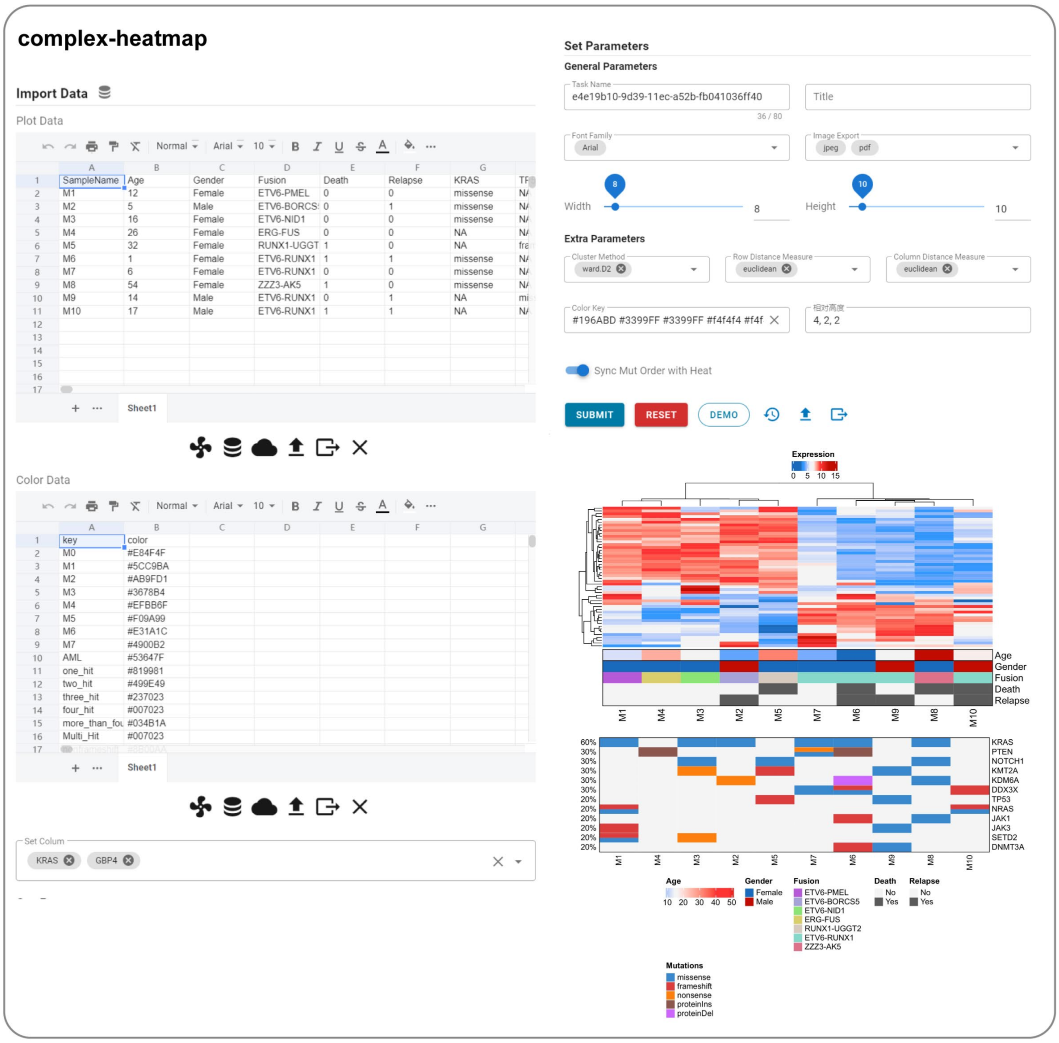


**Fig. S16. Use cases of omics-based data visualizations: multi-omics association analysis based on complex-heatmap plugin.** Web page shows the interface of complex-heatmap. This plugin can be used to conduct the multi-omics correlation analysis based on gene expression data, clinical data, and the genetic mutations.

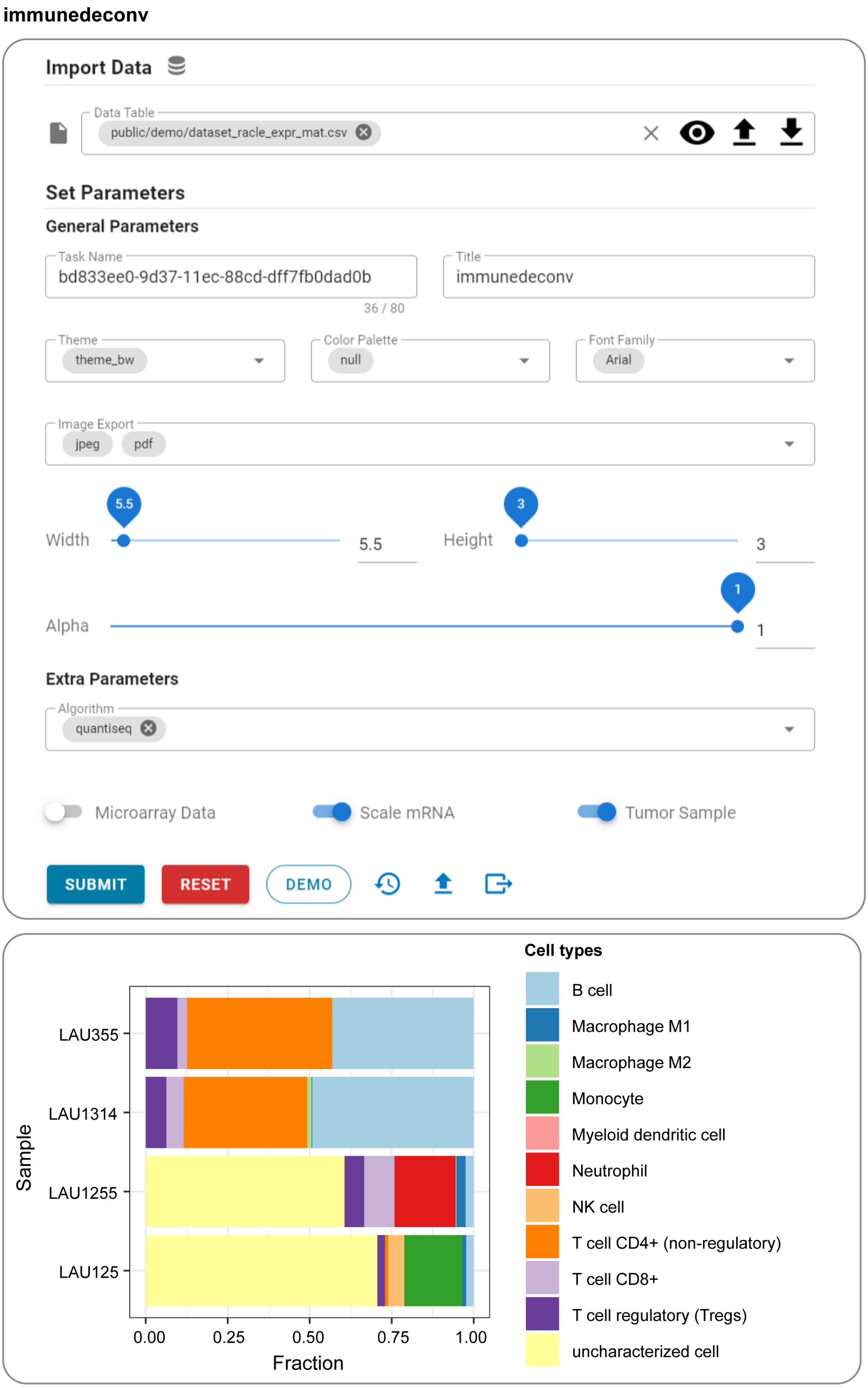


**Fig. S17. Use cases of omics-based data visualizations: tumor micro-environment analysis based on immunedeconv plugin.** Top panel shows the web interface of the immunedeconv plugin, which can be used to conduct tumor micro-environment analysis based on the public algorithms. The bottom panel indicates the bar plot of immune cells fractions in demo samples. Different cell types are filled by different colors.

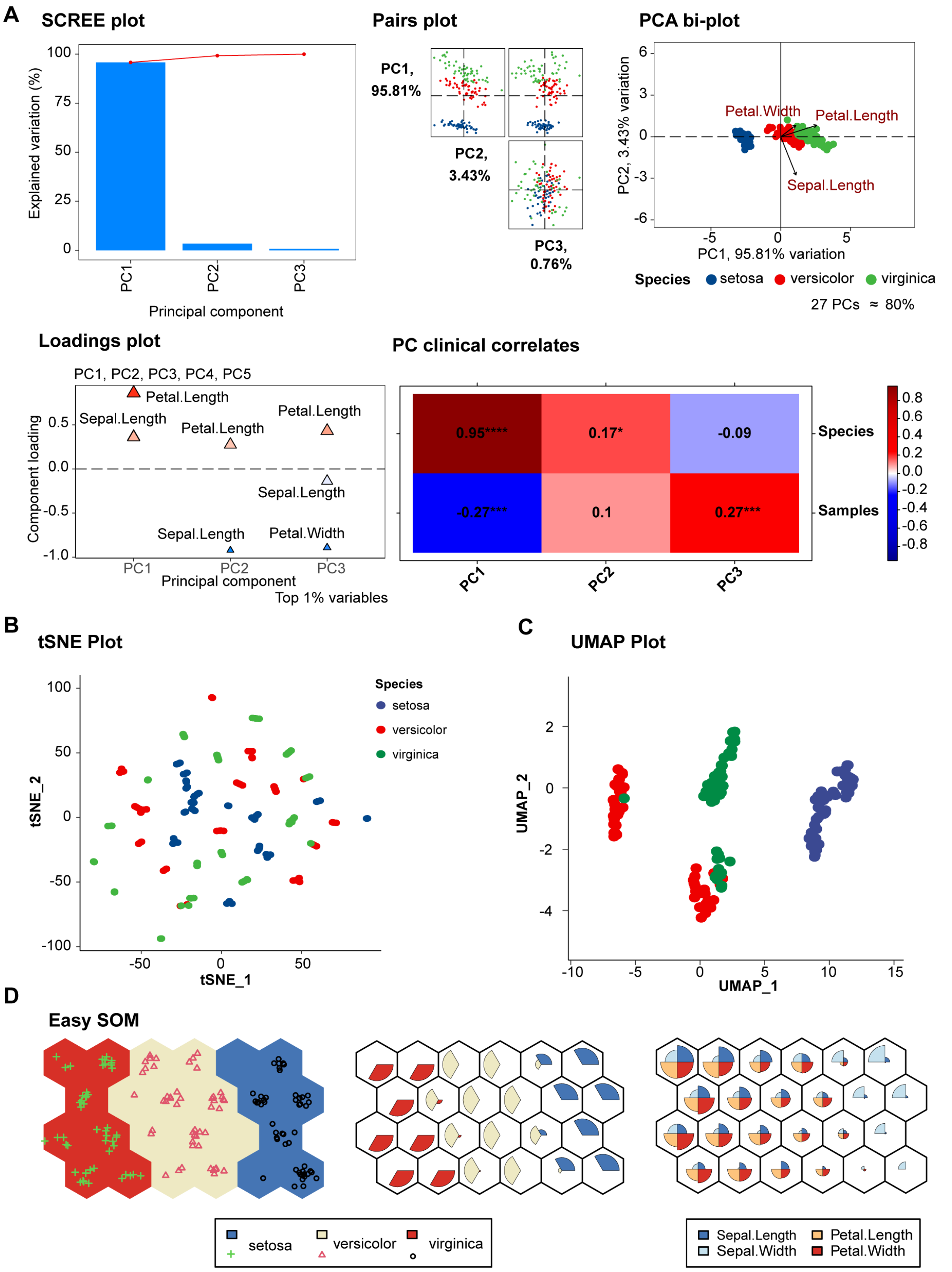


**Fig S18. Use cases of dimensional reduction algorithms.** Demo outputs of (A) pcatools, (B) tsne, (C) umap, and (D) easy-som plugins based on the iris dataset.
